# Heterogeneity of network and coding states in CA1

**DOI:** 10.1101/2021.12.22.473863

**Authors:** Matteo Guardamagna, Federico Stella, Francesco P. Battaglia

**Affiliations:** Donders Institute for Brain, Cognition and Behaviour, Radboud University, Nijmegen, the Netherlands

## Abstract

Theta sequences and phase precession shape the activity time course of hippocampal place cells. Theta sequences are rapid sweeps of spikes from multiple cells, tracing trajectories from past to future. Phase precession is the correlation between a place cell’s theta firing phase and animal position. Here we present an analysis of the strongly variable character of these features in CA1. We identify a cell group that do not phase precess but reliably express theta sequences. The cells that do phase precess only do so when the medium gamma oscillations (60-90 Hz, linked to Entorhinal inputs) dominates but then they show less theta sequences. The same cells express more sequences but not precession when slow gamma (20-45 Hz, linked to CA3 inputs) dominates. Moreover, sequences occur independently in the two groups. Our results challenge a causal relationship between precession and sequences and highlights the highly heterogeneous character of the hippocampal output.

## Introduction

According to current theories [39, 6, 20], the hippocampus combines heterogeneous information from both memory and the outside world to generate representations of episodes and complex occurrences, with space and time as the main organizing principles. Within the hippocampus, the CA1 subfield sits at the confluence of two information streams: the first, directly arising from Layer III pyramidal cells in the Entorhinal Cortex (EC3), carries signals about the animal’s position in space, and highly processed sensory information about landmarks and other relevant environmental features [61, 45, 14]. The second, arising from the Schaffer collaterals coming from the CA3 subfield, is thought to mostly reflect activity shaped by the recurrent connectivity in that area, conducive to memory retrieval and sequence generation [54]. In CA1, these two streams come in contact with each other, providing a neural mechanism that may enable the comparation of memories with incoming external information, and their potential updating [59, 18].

To subserve such a complex function to the rest of the brain, the hippocampus likely makes use of temporal codes. The activity of hippocampal neural ensembles is organized in sequences that can be replayed during short pauses in behavior or sleep [36, 32, 13, 47, 65]. Neuronal sequences also emerge at a compressed timescale in individual cycles of the ongoing theta rhythm, mirroring the order of activation at the behavioral timescale [15, 25]. Since the timescale at which theta sequences occur is compatible with spike-timing dependent synaptic plasticity rules [38], they have been deemed the main mechanism for storing spatial and non-spatial episodic memory traces [17].

Theta phase precession, the systematic backing up in the firing phase of place cells with respect to theta, as the animal advances in the place field [46], has been seen as a key organizing principle for hippocampal temporal codes. While the mechanisms giving rise to phase precession are still widely debated [55, 46, 40, 30, 62, 9], two functional roles have been generally assumed for it. First, phase precession would induce a phase coding [31], that is, the theta firing phase of place cells would be a key conduit for information about the animal’s location. Second, phase precession would be the main ‘engine’ behind sequence formation: under certain assumptions, the presence of phase precession implies theta sequences, that is, a millisecond-scale temporal ordering between the firing of place cells with neighboring place fields, favoring spike-timing dependent plasticity [53, 48, 8]. While theta is most evident in rodent, similar mechanisms may exist also in species where no theta oscillations are observed [21].

Both of these functional roles, however, depend on phase precession and theta sequences taking place in a reliable, nearly noise-free fashion, coherently across neurons, and would break down under high variance conditions. Yet, while we know that phase precession patterns change on a lap-by-lap basis [40, 49], and that theta sequences and phase precession not always co-exist [23, 41, 16], a full picture of how hippocampal temporal coding varies under the influence of the network state is not available. Here, we present a comprehensive analysis of the variability of temporal coding in mouse CA1 place cells. We observe that from being a stable property of all place cells, phase precession varied in degree from neuron to neuron. While some cells showed full-blown phase precession pattern, others were phase-locked to the theta rhythm in a place-independent manner. Furthermore, each neuron could change its phase precessing pattern from one spatial location to the next. On a moment-by-moment basis, “phase-precessing” and “phase-locking” neurons appeared to form distinct functional networks, as they independently express theta sequences in different theta cycles.

The emerging picture suggests that temporal coding depends on which network state and which constellation of inputs each neuron is subject to at any given time. We characterize the CA1 network state by a precise, layer-resolved determination of the instantaneous balance of activity of the medium and slow gamma circuits, which changes from one theta cycle to the next. This is also a proxy for the instantaneous balance of CA1 inputs: inputs from Entorhinal Cortex LIII and CA3 to CA1 respectively are linked, with the emergence medium gamma (60-90 Hz) oscillations in the stratum lacunosum moleculare (slm), whereas the CA3 contribution is associated with slow gamma oscillations (20-45 Hz) in the stratum radiatum (sr) [5, 11, 10, 34, 24]. Medium gamma has been previously associated with stronger phase precession [2] and slow gamma to more precise theta sequences [64]. Here, we show that indeed these two coding schemes are dissociated, and the inputs favoring phase precession are detrimental for theta sequences.

Our results highlight the strong heterogeneity of hippocampal temporal codes, which should be taken into account by any theory of hippocampal function, including those linking phase precession and theta sequence and those assuming that CA1 is generating a unitary and coherent representation through time. The outcoming picture points to a multiplicity of mechanisms and coding schemes being “dynamically multiplexed” in CA1.

## Results

### The Hybrid Drive enables large scale place cell recordings with layer-resolved oscillations in freely moving mice

We implanted 6 mice with the Hybrid Drive (Figure 1A; [28]), a novel recording device that combines linear silicon probes with high density tetrode arrays. Tetrodes were individually lowered to target pyramidal cell bodies in the stratum pyramidale of the CA1 region (Figure 1B,C) and enabled us to collect large ensemble activity of pyramidal neurons (up to *≈* 100) for up to 10 days. The vertical span (960 *µ*m) of the silicon probe yielded LFPs from all CA1 layers (Figure 1C,G and Figure S1).

**Figure 1:**
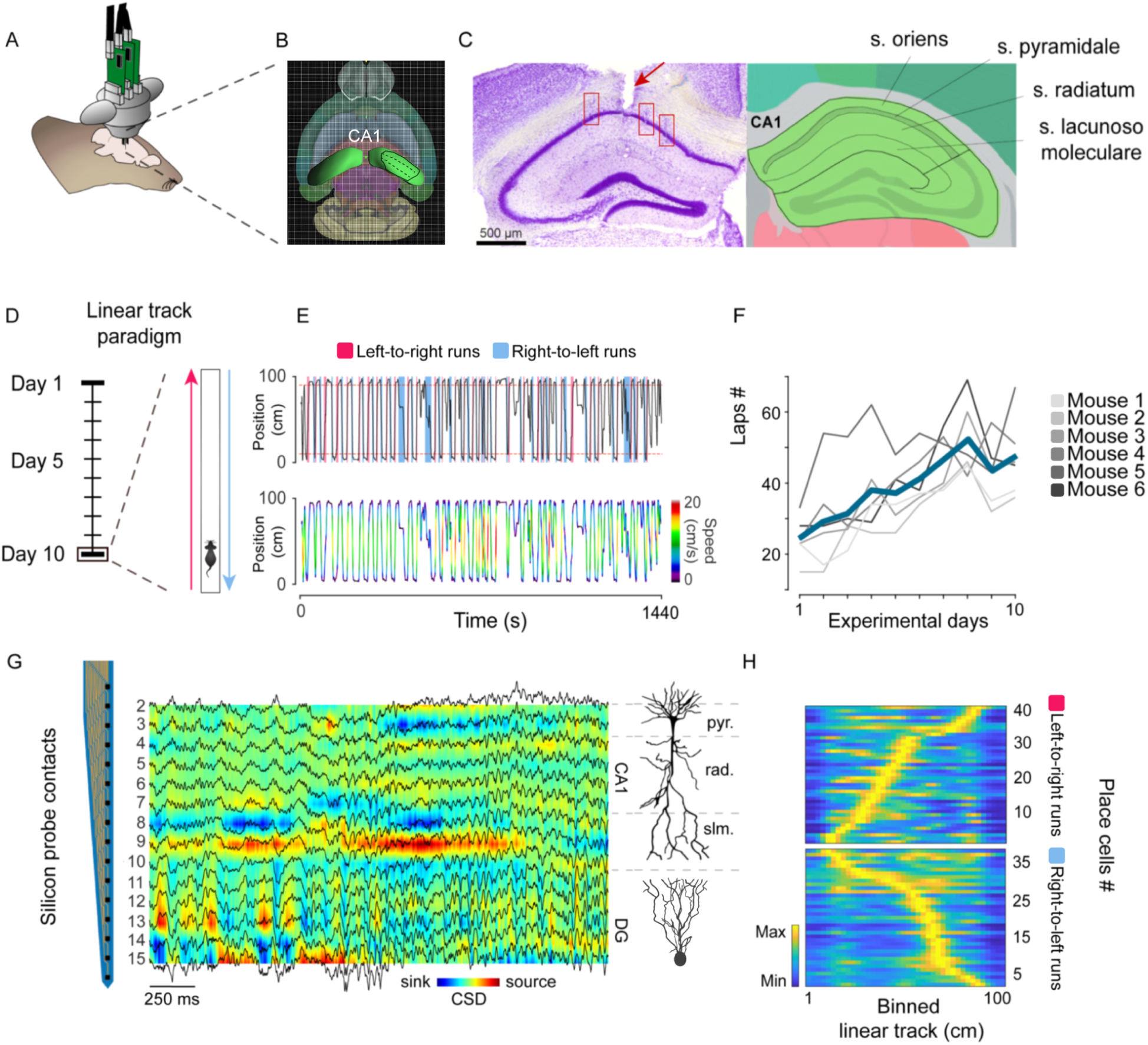
Simultaneous recordings of CA1 place cells and layer resolved oscillations during goal oriented behavior in freely-moving mice. (A) Illustration of the Hybrid Drive implanted on a mouse. (B) Image from the Allen Brain Explorer (beta), a 3D volumetric reference atlas of the mouse brain. Image credit: Allen Institute. The dorsal CA1 region of the hippocampus is highlighted in green. The bold black lines estimate the coverage of the implant array across the distal, intermediate and proximal sub-regions of CA1 . (C) Left: representative histology image from an example animal (Mouse 6). Red rectangles highlighted the tetrode tracks, reaching the pyramidal layer. The red arrow points to the silicon probe entry point. Right: Adapted image from the 2D coronal reference atlas, all CA1 layers. Image credit: Allen Institute. (D) Schematics of the linear track paradigm. 10 consecutive days of training without food or water deprivation. (E) Example of a linear track session from late training days (Day 10). Top: directions of motion are highlighted in two different colors (light pink: up; light blue: down). Bottom: velocity profiles are superimposed on the each lap of the session. (F) Behavioral performance of all animals across the 10 days of linear track training. Average performance is highlighted in dark blue. (G) Color coded depth profile of current sink and sources. LFP traces are superimposed in black. Silicon probe contacts (indicated on the left) span all CA1 layers and partially reach the Molecular Layer (ML) and the Granular Cell Layer (GCL) of the Dentate Gyrus (DG). (H) Example place field maps (same session as in Figure 1G). Each row represents the spatial firing of one cell along the length of the track. Firing rate in color coded and scaled to the peak firing rate for each cell. Cells are sorted according to the position of peak spatial firing on the linear track. Place field maps for both running directions are showed.

This combination of within layer and across-layer recording techniques enabled us to obtain a layer-resolved picture of hippocampal local field potentials (LFPs) (Figure 1G; [34]), while simultaneously monitoring the activity of large populations of place cells (Figure 1H; [35] ) as animals alternated runs between the two ends of a linear track (Figure 1D).

Over the course of 10 consecutive days, mice learned to collect rewards at both ends of a 1 meter-long linear track. While mice were never food or water deprived, we nevertheless observed that performance quickly increased in the first few days and then improved steadily, as measured by the number of laps completed on each day (Figure 1F). For analysis, we considered data from the two running directions separately, by dividing left-to-right from right-to-left runs (Figure 1E, violet and light blue, respectively). Among pyramidal neurons we selected those with significant spatial modulation (see Place Cell Identification). These place cells exhibited place fields that tiled the whole linear track in both running directions (Figure 1H). To investigate properties of stable, well-learned hippocampal spatial representations, we focused on the data obtained from the later recording sessions (between day 7 and 10 of training), which also were the most consistent in terms of number of laps and velocity profiles.

### Fine-scale temporal organization of place cell activity within theta rhythm

Theta oscillations shape the temporal organization of the firing of hippocampal neurons in many ways. There are two simplest models that may account for variability in the data: first, each neuron will fire preferentially around a certain theta phase, i.e. it is phase locked. Second, place cells will display a phase precession pattern as discussed above (Figure 2A). We found high degrees of variability for these two properties across cells and even, for the same cell, depending on the animal location on the track. Thus, cells showed varying degrees of phase locking and phase precession, and these properties are place dependent.

**Figure 2:**
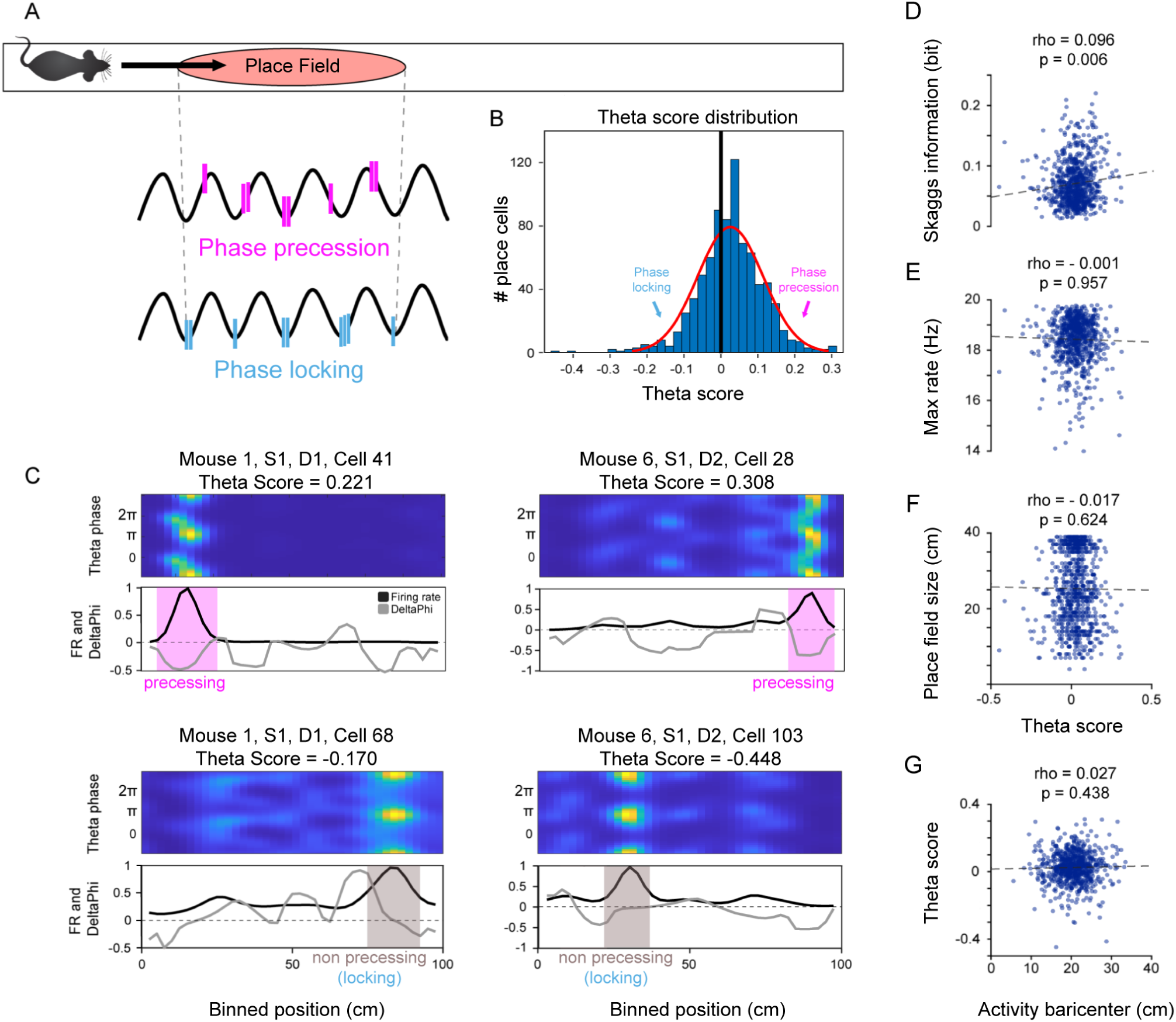
CA1 place cells relationship to the theta rhythm is characterised by a multi-phase nature. (A) Visualization of the two modes of place cells’ temporal organization to theta. Phase precession is described as the systematic advancement of spiking phases as the animal traverses the place field. Phase locking describes the tendency of firing at the same phase of the underlying oscillation. (B) Theta Score distribution of all recorded place cells across all sessions (n=12) and all animals (N=6). (C) Examples from individual place cells, from different portions of the Theta Score distribution. Top to bottom: Color-coded spatiotemporal maps showing the mean firing rate (colour) as a function of position (x) and phase (y); linearized firing rate (FR) maps in solid black lines and phase-position maps (DeltaPhi) in light-gray lines. (D) Scatterplot of Theta Score values versus Skaggs information measure (bits), for field one of all recorded place cells (Spearman’s rho = 0.096; p < 0.01) (E) Scatterplot of Theta Score values versus max firing rate (Hz), for field one of all recorded place cells (Spearman’s rho = - 0.001; p > 0.05). (F) Scatterplot of Theta Score values versus place field size (cm), for field one of all recorded place cells (Spearman’s rho = - 0.017; p > 0.05). (G) Scatterplot of activity baricenter (cm) versus Theta Score, for field one of all recorded place cells (Spearman’s rho = 0.027; p > 0.05).

To fully characterize this variability, for each cell we computed a Theta Score, defined as the difference between circular-linear correlation between theta phase of firing and animal location (a common measure for phase precession [33]) and Rayleigh Vector length (estimate for phase locking). This Theta Score places each place cell on a spectrum and enables us to study how phase locking and precession properties are distributed in populations of place cells. We consistently observed Theta Scores to have an unimodal, Gaussian-like distribution across place cell populations (Figure 2B). Cells at the two ends of the interval presented good approximation of either perfect phase locking or phase precession, but most of the population could be described as a mixture of varying degree of the two modalities (Figure 2B). Theta-score appeared to be largely independent of other properties of place cells: only a very weak correlation was found with the cell Skaggs information (Figure 2D; Spearman’s rho = 0.096; p < 0.01), while no correlation was found with maximum firing rate (Figure 2E; Spearman’s rho = - 0.001; p > 0.05), place field size (Figure 2F; Spearman’s rho = - 0.017; p > 0.05) and place field activity baricenter (Figure 2G; Spearman’s rho = 0.027; p > 0.05 ). Next, we investigated how the locking/precessing properties of a cell vary on the track and how they interact with the mean firing rate at a given location, to provide a picture of the interplay between place cells’ phase and rate coding ([31]). As a location dependent measure of phase precession and locking, we took the spatial derivative of the cell’s preferred firing theta phase along the track (DeltaPhi). As many cells displayed multiple place fields along the track (here, we consider runs in the two running directions separately), by comparing Delta Phi and the location-based average firing rate (FR) we can evaluate the degree of phase precession on a field-by-field basis. In this framework, “phase precessing” fields are described by consistently negative values of DeltaPhi (Figure 2C, top row), whereas “phase locking” place fields display near zero (or positive) DeltaPhi (bottom row examples in Figure 2C). Similarly to the Theta Score, each cell activity relationship with the phase of theta can be summarized by computing the correlation along the track between the cell firing rate and the associated DeltaPhi (phase precessing cells being expected to have an inverse relationship between firing rate increases and negative deflection of the preferred phase of spikes). This measure and the Theta Score are highly correlated (Figure S7A; Spearman’s rho = -0.175; p<0.05).

### Place cells phase coding properties vary across fields

Phase coding modalities are not an intrinsic property of place cells, but rather have a high spatial dependence. In fact, when comparing multiple place fields from the same cell, we found that their phase coding profiles were largely independent from one another (Figure 3A). The probability to precess of field one (defined as the field with the highest peak firing rate; see Place Cell Identification) was only weakly (albeit significantly) correlated (Spearman’s rho = 0.154 ; p < 0.05) to the probability to precess of the secondary field of the same cell (Figure 3B; see Figure S2A,B for spike sorting information for the example neurons). Examples from single-cells reveal how the DeltaPhi measure of field one reflects a phase precessing pattern, while the spikes of the secondary field exhibit a non-precessing relationship, and vice-versa (Figure 3C). Similarly, such phase coding granularity was even more strongly expressed when we considered place cells having place fields in both running directions of the track (Figure 3D). The tendency to precess of a field in one running direction was found to be statistically independent from the phase relationship of the field in the other direction (Figure 3E; Spearman’s rho = 0.092; p > 0.05). The propensity of single fields to precess can be thus conceived as a map-specific property, not unlike the peak firing rate and place field size [22].

**Figure 3:**
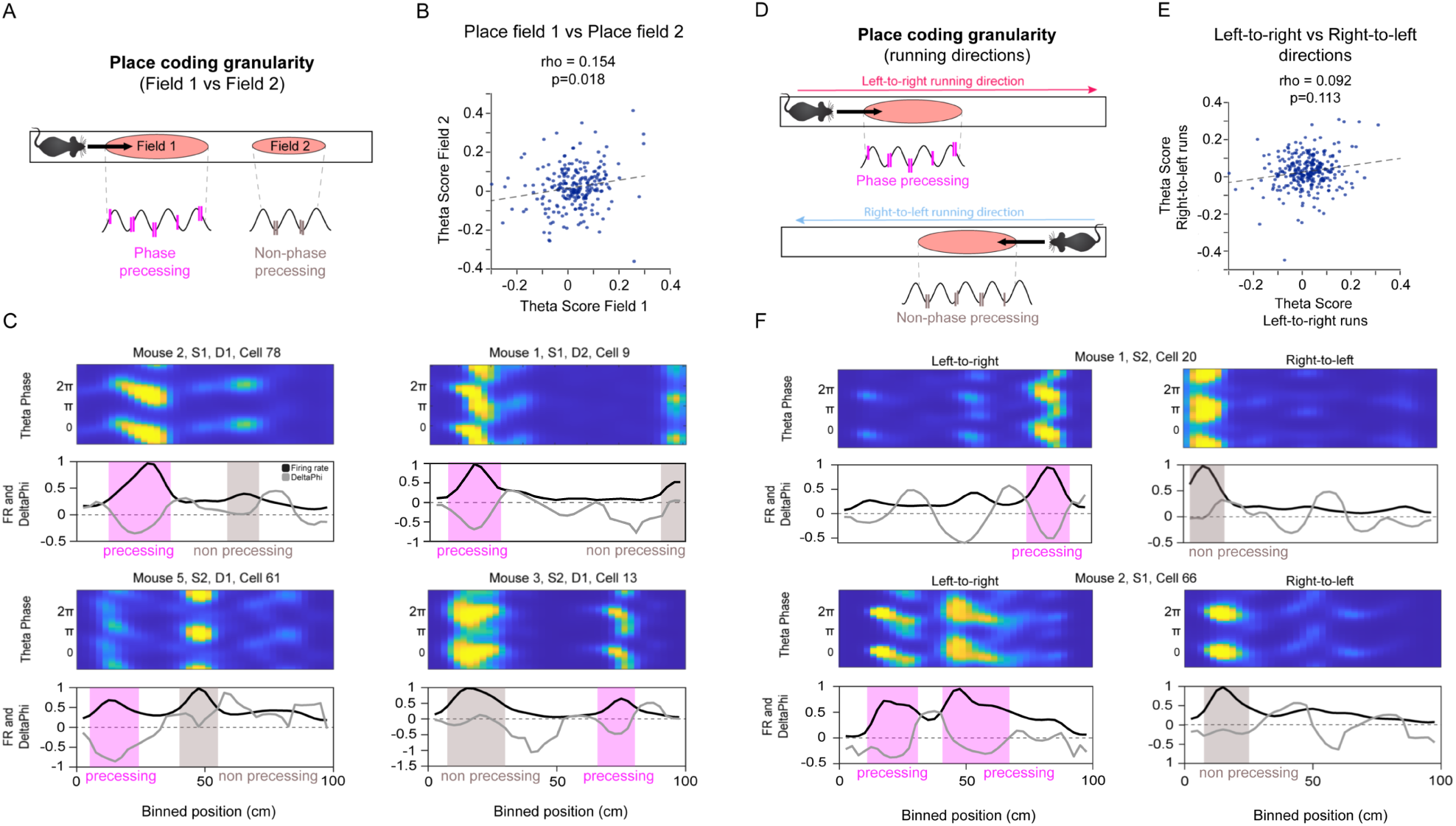
Phase coding granularity between fields and running directions. (A) Schematic representation of independent phase codes (phase coding granularity) in field one and field two. (B) Scatterplot of Theta Score values of field one versus field two, for all recorded place cells with at least two fields in the same running direction (Spearman’s rho = 0.154; p > 0.01). (C) Examples from individual place cells. Note how field one expresses a phase precession pattern while the spikes of field two have a non-phase precessing relationship (top rows) and vice-versa (bottom rows). (D) Schematic representation of independent phase codes (phase coding granularity) in the place fields in the two running directions. (E) Scatterplot of Theta Score values of the field in one running direction (left-to-right) versus the field in the other running direction (right-to-left), for all recorded place cells with two place fields in the both running direction (Spearman’s rho = 0.092; p > 0.05). (F) Examples from individual place cells. Note how the same place cell is expressing a phase precession pattern in one running direction, while in-field phase precession is absent in the other running direction (same session).

### Current source density analysis reveal layer specific theta-gamma interaction in dorsal CA1

After characterizing the spatial modulation of phase precession within CA1 place fields, we sought to explore temporal modulation to phase precession by disentangling the contributions of different input streams in the CA1 circuit, their local integration . In order to obtain a clear picture of the interactions of theta and gamma oscillations in the network we applied Current Source Density analysis (CSD), a standard tool to estimate the contribution of local circuits to the LFP signal and enables precise layer-level resolution (which, in the hippocampal CA1 circuit relates neatly to different afferent pathways and groups of cells). For each CSD-time series we then computed the phase-amplitude relationship between the theta oscillation (6-10 Hz) and the power of oscillations at higher frequencies (15-250 Hz). Since theta power is maximal in the deeper portions of CA1 and theta oscillation presents a high degree of coherence across layers, we decided to use the signal from the slm probe contact as the reference theta oscillation signal. This signal was then used to produce a depth-specific profile of frequency-resolved gamma-theta couplings. Note that slm theta is 180 degrees shifted from pyramidal layer theta, used as a reference in some studies.

Following Lasztóczi and Klausberger [34] we characterized gamma oscillations in a tridimensional ’gamma space’ with anatomical depth, frequency, and preferred theta phase as dimensions. Significant components in this space were then defined as contiguous regions of above-mean correlation strength. Once points in gamma space with significant coupling values found at different depths were concatenated, we obtained a 3-dimensional structure that represents theta-gamma interactions (Figure 4A). We then isolated individual volumes of significant coupling with consistent spatial extension and termed them ’Basins’. These Basins provide a data-driven definition of independent gamma oscillators in the CA1 circuit. We found 3 of these Basins, coinciding with 3 separate ranges of gamma: 20-45Hz, 60-90Hz and 100-150Hz. In the following they are identified as low (Figure 4B), medium (Figure 4C) and fast gamma (Figure 4D), respectively. Each component can be characterized by its distinctive depth profile, most evident in the location of maximal coupling with theta phase (Figure 4F). While slow gamma (Figure 4B) showed minimal variability along the CA1 depth profile, with a mostly constant theta coupling strength across CA1 layers (Figure 4F), we could observe a gradual shift in the preferred coupling theta phase from more superficial layers (pyramidal) to deeper ones (slm) (Figure 4E). On the contrary, the medium gamma component (Figure 4C) displayed a spatially localized pattern, with coupling strength peaking in the slm (Figure 4F). Different from slow gamma, medium gamma theta phase preference shifts abruptly between the pyramidal layer and the stratum radiatum, and stays constant in the deeper layers (Figure 4E). Finally, the fast gamma component (Figure 4D) emerges only in proximity of the pyramidal layer, maintaining a constant phase preference value (Figure 4E).

**Figure 4:**
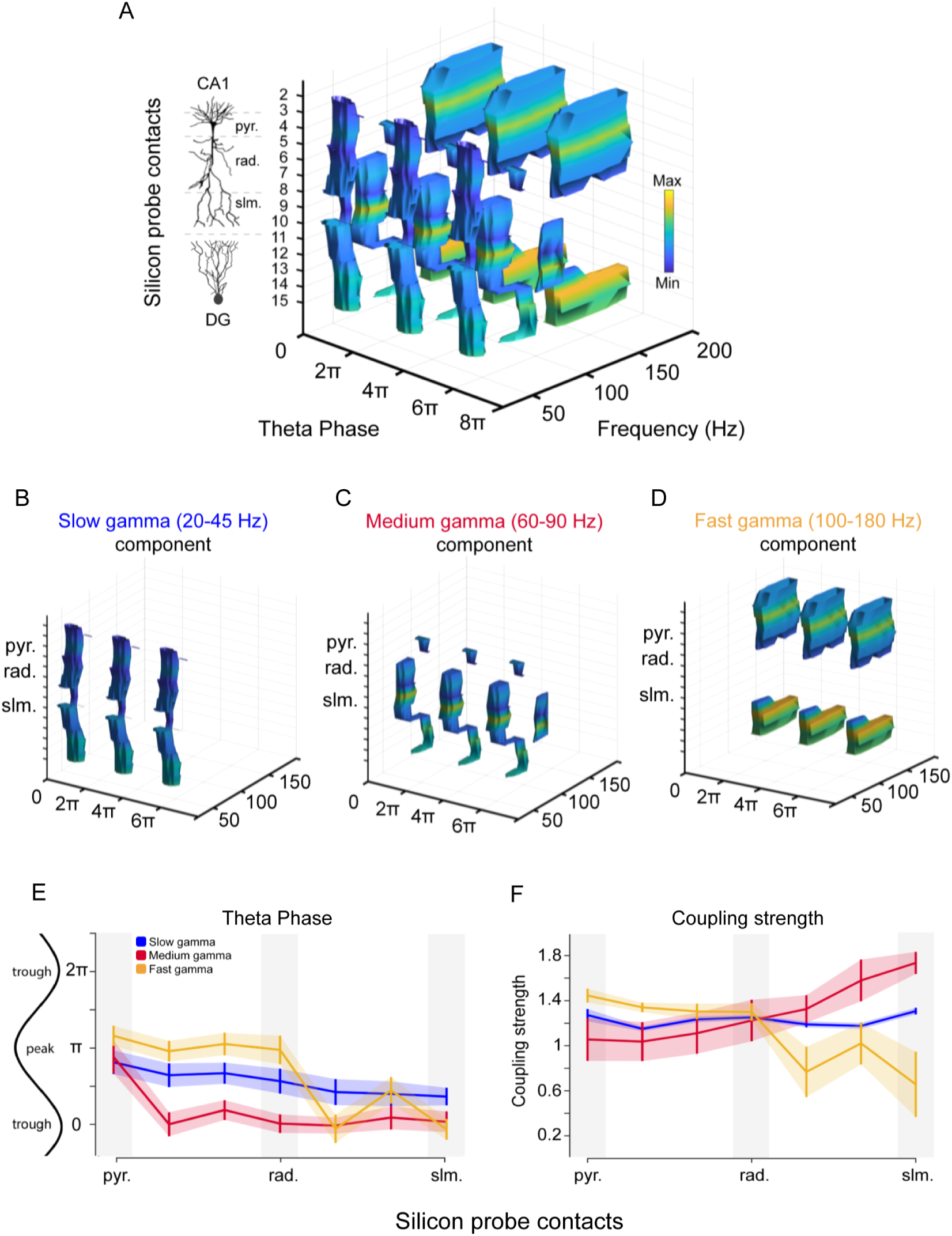
Theta-gamma interactions in a 3D phase-amplitude-depth structure. (A) CSD-derived Basins profiles, along one shank (16 recording sites; spaced at 60 *µ*m) spanning the entire CA1 region (2-10) and part of the Dentate Gyrus (12-15). The Basin structure is repeated across 3 theta cycles. (B) Isolated slow gamma (20-45 Hz) Basins component from panel (A). (C) Isolated medium gamma (60-90 Hz) Basins component from panel (A). (D) Isolated fast gamma (100-180 Hz) Basins component from panel (A). (E) Theta phase preference profiles of the three Basins components across all CA1 layers. (F) Coupling strength profiles of the three Basins components across all CA1 layers.

Based on this detailed description and visualization of gamma generators provided by the Basins, we then set out to disentangle the contributions of these circuit elements to the spike timing of place fields and place cell ensembles.

### Phase precession arises in a subset of place fields when medium gamma dominates the CA1 network state

Because place fields do not appear to have consistent phase coding properties, we speculated that their temporal coding modalities may be affected by variability in the inputs to CA1, reflecting changing environmental conditions. We therefore investigated how the cycle-by-cycle balance of different gamma regimes, considered proxies for the influence of those different input sources [11, 35], control the organization of the phase precession pattern within individual place fields. We used a Generalized Linear Model (GLM) to describe the spike probability of each place field given i) the position within the place field; ii) the instantaneous theta phase and iii) the instantaneous power in a specific gamma range and layer (as they were obtained from the previous Basins analysis). Depending on the Theta Score values of field one (described as the field with the highest peak firing rate; see Place Cell Identification), we divided the place field population in three equally distributed groups. Intermediate fields occupy the center of the Theta Score distribution, while phase precessing and phase locking the extremes. Crucially, we found that fields within the phase precessing population, were not performing phase precession in every theta cycle, but rather that phase precession could be seen to gradually appear in association with the increasing presence of medium gamma in the theta cycle. The spike density plot in Figure 5A reveals how the phase precession pattern gradually emerges as medium gamma becomes more dominant in the CA1 network state. The contour plots of the spike density distributions show how the appearance of spikes in the early portion of the theta cycle is conditioned to the increase of medium gamma power (Figure 5B). Conversely, changes in gamma balance do not affect the spike distributions of the phase locking and intermediate population. This is replicated for each animal separately (Figure S4). Importantly a further analysis using velocity as an additional factor in the GLM, shows that the observed gamma modulation of the phenomenon of phase precession is independent of changes in the running speed of the animals (Figure S6).

**Figure 5:**
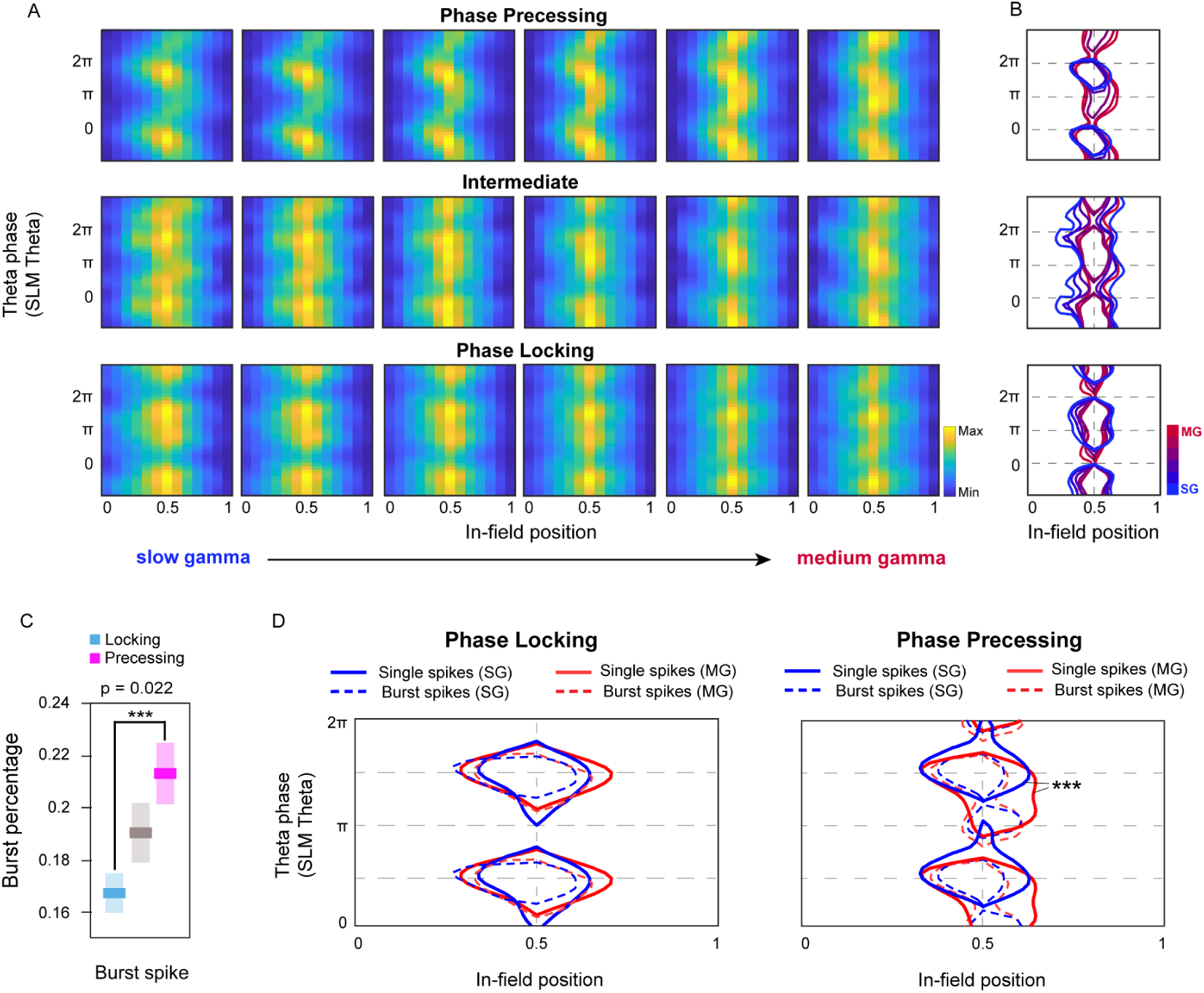
Phase precession arises in a sub-population of place fields during periods of medium gamma dominance. (A) Spike density plot of all cells from one example animal (Mouse 1). Global average of the in-field spike probability, plotted as a function of the instantaneous gamma coefficient, for each subset of place fields (precessing, intermediate, and locking). The phase precession pattern gradually emerges only in the phase precessing subgroup, with increasing medium gamma power. (B) Contour plot of the same Spike density plot in (A), highlighting the progressive, selective emergence of a phase-location dependence for precessing cells (in association with medium gamma dominance). (C) Burst fraction in the phase precessing, intermediate and phase locking populations (p = 0.022, two-sample t-test). (D) Contour of regions with highest density of spikes for single spikes during slow gamma (solid blue line) and medium gamma (solid red line), for burst spikes during slow gamma (dashed blue line) and medium gamma (dashed red line) network states. Kolmogorov-Smirnov test on spike densities (*** = p<0.01).

Long dendritic depolarizations [26, 43, 3] may induce bursts of spikes in CA1 pyramidal cells, and it is therefore interesting to explore whether bursting dynamics differs between precessing and non-precessing fields. We considered separately spikes that where either emitted in bursts or in isolation (ISI>7ms; see Single Spikes and Burst Spikes classification) and we found a significant modulation of burst propensity by Theta Score, with phase precessing fields having on average an higher burst fraction (Figure 5C; p<0.05, two-sample t-test).

Contour of regions of highest density of spikes further reveal how the population of phase precessing fields is more affected by the instantaneous balance of gamma oscillations; while the population of phase locking fields tends not to vary the in-field/phase organization of spikes (Figure 5D; Kolmogorov-Smirnov on spike densities, all animals (N=6), all sessions (n=12)). Moreover, for phase precessing cells the higher amount of burst events was not evenly distributed over the theta cycle. During medium gamma dominated cycles they appeared to concentrate at the earliest phases of the cycle, ahead of the phase interval containing most single spikes; while during slow gamma dominated cycles the phase preference of burst spikes was more evenly distributed along the phases of the ongoing theta cycle (Figure S7C; significant Rayleigh Vector Length Difference against shuffling, T-test: p<0.01).

When looking at single field cases (phase-position slopes of individual fields), the effect is still present. With increasing dominance of medium gamma, fields with a higher Theta Score tend to acquire a steeper phase-position slope (Figure S5). This medium gamma related effect is only visible for fields with higher Theta Score, while slow gamma does not modulate the phase-position slope and the Theta Score. Once again this is replicated for each animal separately (Figure S5). Together, these observations suggest that phase precession selectively arises in a subset of place fields, as a function of the oscillatory state of the CA1 network.

### Organization of theta sequences during slow and medium gamma network states

Until now, we have been looking at phase coding, a potential way for single receptive fields (place fields), to convey spatial information in addition to what is already contained in the firing rates [31]. Yet, a key role that theoretical models ascribed to phase precession is the shaping of the activity of one place field with neighbouring fields into temporal sequences spanning one theta cycle [53]. Because we observed an effect of gamma oscillations on phase precession, we therefore asked whether such theta sequences were likewise affected by changes in medium and slow gamma power. We first addressed pairwise co-activation patterns, a basic, but sensitive index of sequential activation [52, 15, 41]. Similar to [15, 41], we measured the correlation between the distance between the place fields of a place cell pair and the average theta phase difference of their first spikes in each theta cycle (Figure 6A,B). This correlation measure reflects how much a spatial distance could be reliably inferred from a temporal one. We repeated this procedure by further separating spikes according to the gamma dominating the theta cycle in which they were emitted.

**Figure 6:**
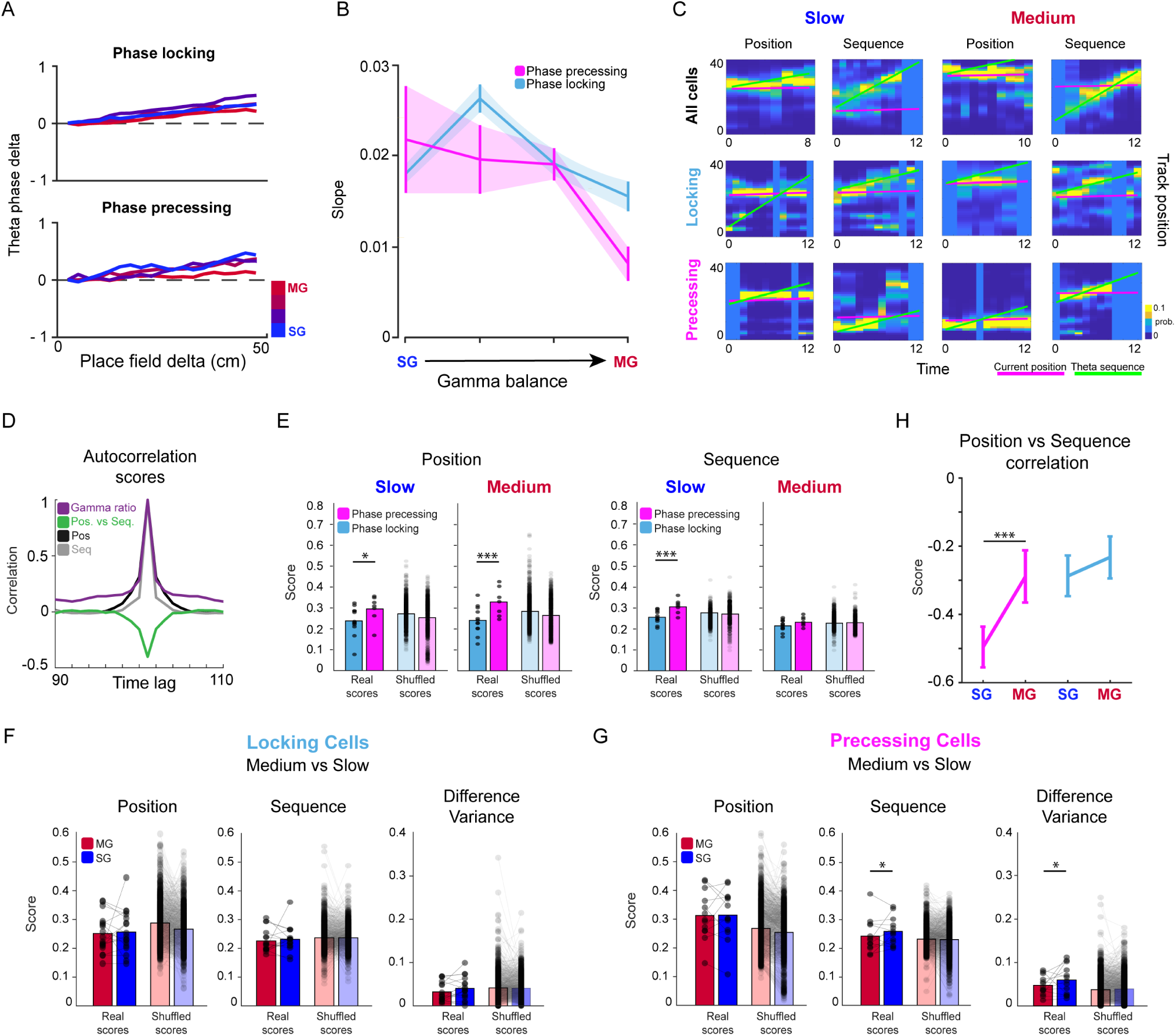
Theta sequences organize during slow gamma dominated network states. (A) Place field distance vs phase distance regression for different gamma coefficients and populations (regression lines were aligned to their first bin on the x-axis). (B) Summary of regression slopes shown in panel (A). p=10*^−^*6, confidence intervals, during medium-gamma network states. (C) Examples of encoded position (y-axis) recon-struction over an entire theta cycle (x-axis), dividing theta cycles according to the corresponding slow-medium gamma power balance and isolating spikes from specific groups of cells (all cells, or phase locking and phase precessing cells). Blue-yellow color scale: posterior probability from Bayesian Decoding. Violet line: real position of the animal, used to compute the amount of local coding as the probability density around the line. Green line: best positive-slope fit of probability distribution, used as an estimation of sequential non-local cell activation. (D) Theta-cycle based temporal cross-correlation of decoding scores and gamma power. (E) Comparison of mean positional and sequential scores between locking and precessing cells in slow and medium gamma conditions. Points: sessions. (shaded bars: control results from shuffling of theta scores across the population of simultaneously recorded cells). T-test (*** = p<0.01; * = p<0.05) (F) Comparison of mean positional and sequential scores and of the variance of their difference between different gamma conditions for cells classified as phase locking. (shaded bars: control results from shuffling of theta scores across the population of simultaneously recorded cells). Paired t-test (* = p<0.05) (G) Same as (F) for cells classified as phase precessing. (H) Zero-lag cross-correlation between positional and sequential coding for different gamma conditions and theta-score cell groups. T-test (*** = p<0.01)

Measuring the gamma-dependent spatio-temporal correlation between spike pairs lead to two results. First, fields characterized by a strong phase locking showed a significant degree of spike ordering, independent of the gamma configuration at the time of the spike emission. Second, phase precessing fields, on the contrary, presented a variable amount of temporal ordering, depending on the slow-medium gamma balance. Crucially, spatio-temporal correlation of spike pairs was only present in coincidence with strong slow gamma periods, that is, only at times when phase precession was at its weakest (Figure 6A,B). Therefore the ability to carry information about macro-scale ordering of place fields within the micro-scale arrangement of spikes at the tens of milliseconds scale differ across fields, and depends on the current state of the networks generating gamma in CA1. Importantly, the states of these gamma networks conducive to phase precession and to pairwise ordering of spikes appeared to be complementary to one another. Pairwise interactions between fields provided a first approximation to evaluate the degree of which neural activity in CA1 place cells populations is sequentially organized. We took this analysis a step further by implementing a decoding procedure, aimed at estimating the level of sequential information contained in the entire population of recorded place cells. Because this type of analysis cannot distinguish between spikes belonging to different fields of the same place cell, we extended our definition of ’locking’ and ’precessing’ to cells rather than single fields. Each cell was assigned the identity of its place field with the highest firing rate (Field 1 as shown in Figure 3A, when multiple fields were present). By dividing each theta cycle into a set of small sub-windows, we computed the likelihood of the activity contained in each of them to encode one of the positions of the track. Using these probability density matrices, we evaluated the degree of non-locality of the encoded positions, that is to what degree the encoded positions within a theta cycle a) reflected the current position of the mouse on the track or b) formed a “theta sequence”, that is, they represented a set of positions sequentially distributed in time around and independently from the current position of the animal. Again, such computation was repeated separately using spikes from either the entire population, or from specific groups of cells and also by subdividing theta cycles according to the corresponding slow-medium gamma power balance. Results obtained with Bayesian decoding confirmed and expanded those based on pairwise measures (Figure 6C). We first considered the temporal organization of the positional and sequential coding scores by computing the auto-correlation of the scores obtained for theta periods at different time lags. The encoding of sequences showed almost no temporal correlation already in neighbouring theta cycles, while positional coding appeared to retain some degree of continuity within short stretches of theta cycles (Figure 6D). As partially expected by their definition, the two scores turned out to be anti-correlated within the same theta cycle. The spatial information content carried by either locking and precessing cells was found to be different and to be gamma-modulated (similarly to what found with pairwise ordering): while on average precessing cells better encoded position, the representation of spatial sequences over the course of a theta cycle was selectively enhanced in precessing cells during periods of strong slow gamma power. During periods dominated by medium gamma instead the sequential coding was comparable in the two populations of cells (Figure 6E). Indeed when we directly measured the effects of relative gamma balance in each of the two populations, we found no effect in the locking group and an enhancement of sequential coding for precessing cells (Figure 6F,G). Interestingly such increase in sequential-like activation in precessing cells led to a widening of the position-to-sequence score range (computed as a difference between the two scores). That is, during slow gamma periods each theta cycle was more strongly biased towards either positional or sequential coding. We directly measured this temporal segregation of spatial-related coding schemes by measuring the cycle-by-cycle correlation of the two scores. On top of the previously noted anti-correlation, we found a gamma-induced modulation in precessing cells. Consistently with the previous measure of the score difference variance, the anti-correlation between the two scores was found to be stronger during periods of slow gamma prevalence (Figure 6H). Consistently with our previous analysis, thus, the population approach confirmed that for specific subsets of cells phase precession and sequential coding emerged in CA1 under different network conditions.

### Independent spatial coding in hippocampal sub-populations

We then asked how the instantaneous expression of a specific spatial coding scheme coordinated across the CA1 network. To do so we first computed again the pairwise spatio-temporal correlation we used before, but this time considering cell pairs across the two cell groups of locking and precessing fields (Figure 7A). We found that slow gamma modulations reshaped these groups: an increase in the slow gamma power coincided with an increase in the pairwise score for cell pairs within the same theta-score group (thus strengthening sequential coding within each group) and a small but significant decrease in the correlation of across-group pairs (although this effect was stronger for precessing cells), thus decoupling the two groups. When medium gamma dominated, the cell pair slopes calculated within and across groups were indistinguishable, and were also not different from reshuffling the theta scores across cells from the same session. Together, these data suggest that slow gamma increases order in the precessing and in the locking group separately.

**Figure 7:**
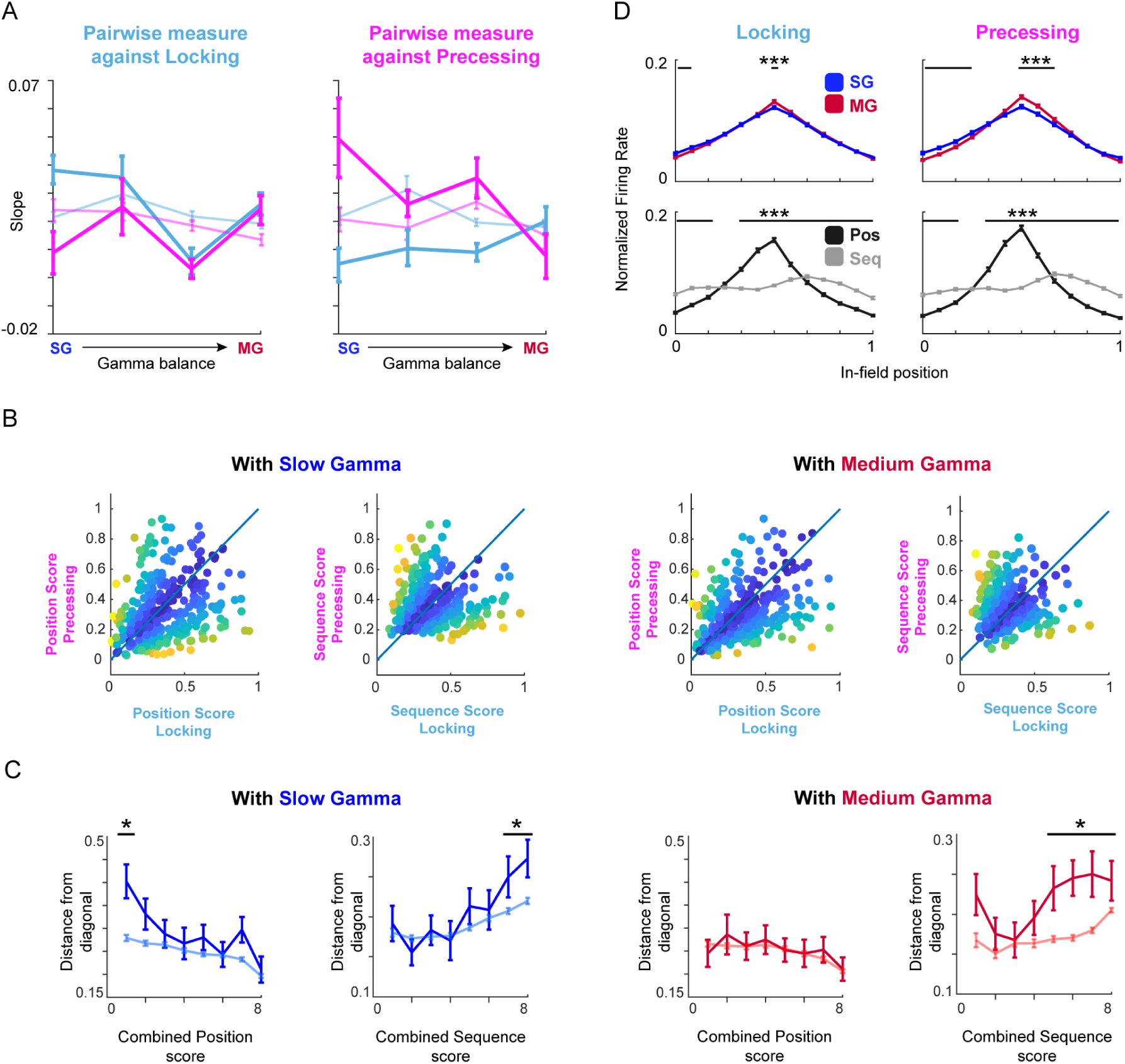
Locking and precessing cells show segregated spatial coding. (A) Spatio-temporal pairwise spike correlation computed for cell pairs either within the same cell group or across theta-score cell groups, as a function of the instantaneous gamma balance. (B) Distribution of simultaneously measured decoding scores (left: position; right: sequence). Each point represents a single theta cycle and its position is defined by the score obtained using either only spikes from precessing cells (y-axis) or only from locking cells (x-axis). Left panel: data from temporal periods dominated by slow gamma. Right panel: data from temporal periods dominated by medium gamma. Color coding represent distance from the diagonal. (C) Average distance of points in (B) from the diagonal (corresponding to a perfectly equivalent coding in locking and precessing cells) as a function of their combined value. Left panel: data from temporal periods dominated by slow gamma. Right panel: data from temporal periods dominated by medium gamma. Shaded lines represent shuffled distributions. T-test (* = p<0.05). (D) Place field shape obtained when considering different temporal intervals. Top: slow vs. medium gamma dominated theta cycles. Bottom: temporal windows defined by the type of spatial coding observed by selecting theta cycles based on the position vs. sequential decoding scores. T-test (*** = p<0.01).

We observed a similar phenomenon when considering the results of Bayesian decoding. In this case we compared the decoding scores obtained from different place cell groups for the same theta cycle. Inspecting the distribution of this simultaneous decoding score combining locking and precessing cells (Figure 7B) we observed that while for positional coding points tended to concentrate around the diagonal (indicating a high degree of coordination between the coding in the two group of cells), in the case of sequential coding points tended instead to be located away from the diagonal, especially for increasing values of the score, showing that the two groups express theta sequences in separate theta sequences and rarely simultaneously. We quantified this effect by measuring the average distance of points from the diagonal as a function of the combined decoding score for each theta cycle (Figure 7C; see Methods). This analysis confirmed our previous observation, showing a significant distancing of high-score sequential events from the diagonal and a substantial uniformity in the case of positional coding.

Finally we compared the place field shape in the different temporal windows defined in our analysis. First, the effect of gamma modulation was identified in a backward shift of place fields in correspondence with periods of high slow gamma power, with the effect being stronger for precessing fields (Figure 7D, top panels). Then we also observed that identifying temporal windows based on the nature of the expressed spatial coding (either positional or sequential) led to a measurable change in the shape of place fields. In fact, when taking only periods characterized by an high degree of sequential organization of spikes resulted in the flattening of the measured place field shape, with no observable rate tuning left (Figure 7D, bottom panels).

Interestingly, further evidence for the presence of a theta-score based segregation of cell groups organizing in theta sequences was given by looking at the enlargement of fields when detecting sequences separately for locking or precessing cells (Figure S8A). Indeed we could observe a double dissociation: periods of precessing cell detected sequences corresponded to an enlargement of precessing cell fields, but to no significant modulation in the locking cell fields, and vice versa.

## Discussion

The data we presented here highlights the heterogeneity of temporal coding schemes in CA1 activity. We first showed that phase precession is not a general feature of place fields (Figure 2). Rather, we observed a continuum between purely phase locked and phase precessing fields, with no or very little correlation with spatial tuning and neural activity parameters. Fields at the “precessing” and “locked” ends of this spectrum differ in non space-related properties, such as their theta phase locking during active behavior (or during REM sleep; Figure S7B), suggesting that intrinsic differences in anatomy and physiology, as for example belonging to the deep or superficial pyramidal sublayer [42, 57, 58, 50] may affect only to some extent a place field’s propensity to phase precess. Yet, by analyzing phase precession as a function of location on the track (Figure 3) we show that the same place cell may substantially vary its phase precessing behavior at different locations on the track. Thus, intrinsic cell properties may explain only a small portion of the total variance at best, and the causes of phase precession may have to be searched not only in the circuit layout, but, most poignantly, in the precise constellation of inputs, cellular and network states affecting a neuron in a given contingency.

Second, when looking on a theta cycle-by cycle basis, CA1 population activity either expresses a forward-sweeping (on average, but see [60]) sequence, or a stable representation of animal location. The sequential content of the representation fluctuates very rapidly, and the probabilities to observe theta sequences in consecutive theta cycles are virtually uncorrelated (Figure 6D). Third, theta sequences are not expression of the CA1 cell population as a whole. Rather, locking and precessing subpopulations may form, at least momentarily, separate coherent groups, as shown by the fact that they express theta sequences independently and at different times (Figure 7). For example, it is possible that one group expresses a sequence while the other encodes current position. Whether a single “position readout” decoded from CA1 activity is fully representative of information processing in the hippocampus is called into question by these results.

Fourth, rather than being one the cause of the other, phase precession and theta sequences are by-and-large independent, if not downright in opposition, meaning that they tend to occur at different moments, at least in one group of cells. Our results point at different CA1 inputs being responsible for the two phenomena, with CA3 inputs being conducive to sequences and Entorhinal LIII inputs to precession. It has been previously shown that strong slow gamma periods (putatively, with predominant CA3 influence) are related to tighter theta sequences [64]. We show here that the effect is to be ascribed to phase precessing fields only, whereas cells that do not exhibit phase precession reliably show sequences regardless of gamma oscillations (Figure 6). Conversely, increased medium gamma intensity (proxy for EC LIII inputs) is related to tighter phase precession [2]. Here as well, we show that this is only due to the phase precessing fields, whereas phase locking cells never precess. Furthermore, we characterize how the phase precession pattern arises as a new cluster of spikes emerges in the position by firing phase plane (Figures 5 and 8), at the trough of the slm theta. That cluster of spikes may act as a disturbance for theta sequence expression, causing the decline in sequence score displayed in Figure 6.

**Figure 8:**
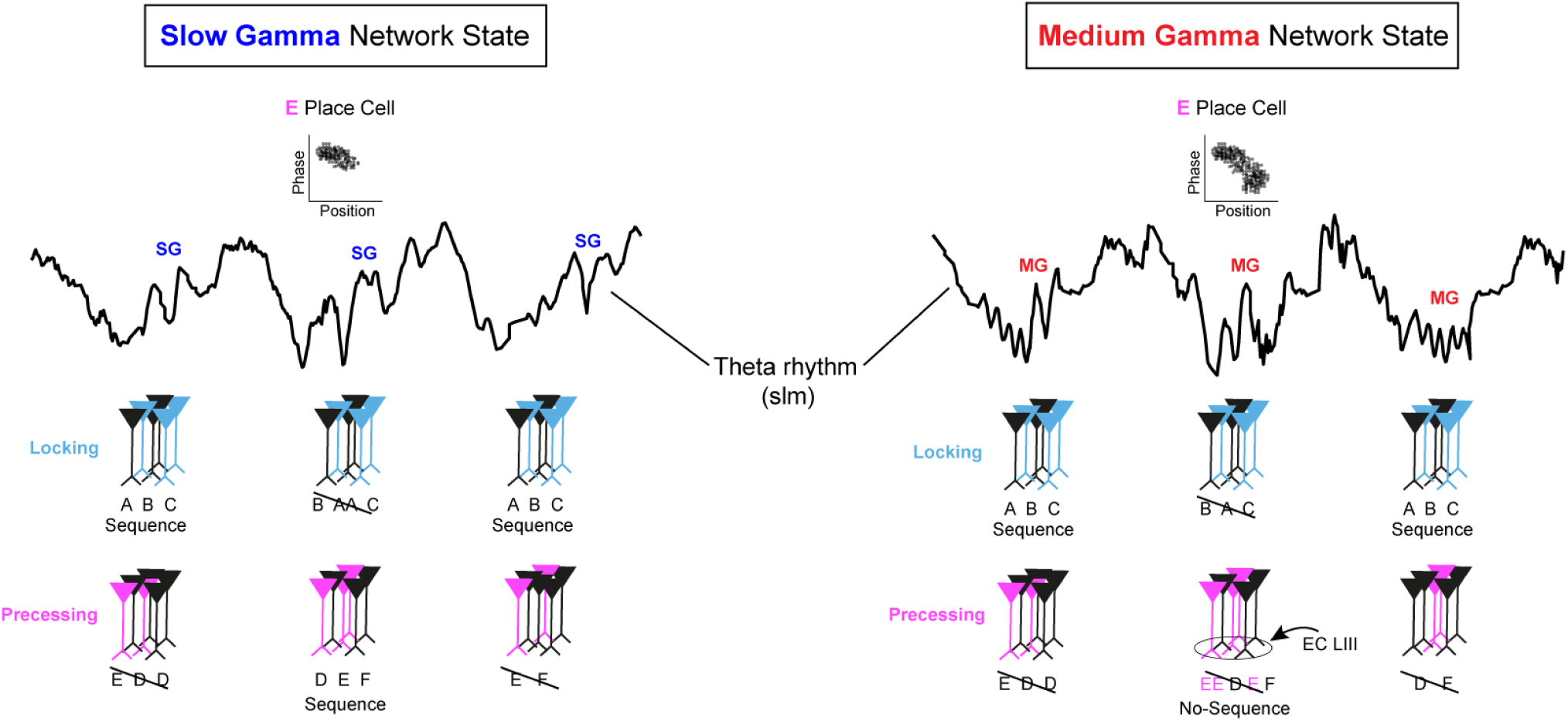
A comprehensive framework for CA1’s heterogeneous temporal coding dynamics. We summarize our observations and previous literature by assuming that the CA1 network spontaneously fluctuates between two dynamical states, one with strong CA3 inputs and slow gamma oscillations and one dominated by EC LIII inputs and medium gamma. Furthermore, we assume two groups of neurons, “locking” and “precessing”, which form theta sequences independently and at different times. Similarly, phase precession is not expressed homogeneously by all CA1 place cells, but is modulated as a function of space and time (Figure 3 and Figure 5). When slow gamma dominates (left), cells that should phase precess at that location only fire at the theta phase of maximal CA3 inputs. Spiking takes place in the initial portion of the place field, so that spatial tuning is prospective. Both phase locked and phase precessing cells display theta sequences (independently) in this state. The full phase precession pattern appears in the “precessing” subset of cells when medium gamma is strongest (right). For those cells, a second cluster of spikes appears on the position-phase plane, shifted towards an earlier theta phase, corresponding to EC LIII activation and to a further position in the place field. The relative position of the two clusters gives rise to the phase precession effect [9]. Bursts are most prominent in the second cluster, suggesting the influence of dendritic dynamics triggered by EC LIII afferents depolarizing distal apical dendrites of pyramidal cells. The interference between spikes in the two clusters disrupts theta sequences, which are still apparent in phase locked cells. This state may favor the combination of memory-derived and new sensory information.

The phase precession phenomenon [46, 30, 40, 55] and its links to theta sequences [53, 8, 48] have been the object of multiple theories and models. We believe that high level of heterogeneity and temporal variability shown here represents a challenge for all those mechanistic theories. Instead, a simpler account by F. Chance [9] fit the present data quite well. In this model – as already hypothesized by Yamaguchi et al. [62]) – spikes form two clusters in the animal position by theta phase plane. The cluster corresponding to entry into the place field and later phase is ascribed to CA3 inputs (Figure 8). EC3 inputs would be instead responsible for the spike cluster at the exit of the place field. The theta phases of the two clusters is consistent with CA3 inputs lagging EC3 inputs by a quarter theta cycle. If we assume that medium gamma covaries with the EC3 inputs strength, that model would explain the result in Figure 5 (and in [2]. The Chance model, augmented with time-fluctuating CA3 and EC3 input strengths, and slow and medium gamma intensity being their respective proxies, is consistent with a number of observations in the literature. In particular, the model would predict the fact that slow gamma induces a backward (prospective coding [1]) shift in place field location [2, 7] whereas the field moves forward (retrospective coding) with elevated medium gamma. We reproduced this effect in our data (Figure 7D). Furthermore, spike-triggered averages of medium and slow gamma power respectively increase and decrease with place field traversal [34] (also reproduced here Figure S8B).

As an important technical aside, here we recorded laminarly resolved LFP and neural ensembles are recorded from separate sites [28] (Figure 1), which avoids any spillovers of spiking activity in the LFP, which may bias LFP-spikes association measures. We then used a novel analytical procedure (Figure 4 and S3), derived from ref. [34] to separate the different generators at different hippocampal layers, making explicit use of preferred theta phase information. It is possible that some discrepant results in the literature [24] are due to different approaches in computing the time series of gamma intensity .

In sum, we show here that there is a cell population, hereby named “phase precessing”, that shows a broader range of temporal coding regimes, and changes most markedly as the input balance in CA1 changes, with respect to the rest of CA1 neurons. The makeup of this group varies even from one spatial location to the next. An enticing hypothesis is that the distribution of inputs on the dendritic tree is the determinant of this behavior. Dendritic targeting interneurons may shift the balance of the inputs between CA3 and EC3 [37, 56, 27]. It is also known that EC3 inputs induce “plateau potentials” in CA1 pyramids’ distal apical dendrites [3, 44, 4] and that these may last for hundreds of milliseconds, possibly linking inputs occurring within that time interval. The testable prediction is that cells that would be classified here as phase precessing are those that are mostly activated by dendritic branches under the tightest interneuronal control and/or expressing more plateau potentials. Which dendritic branches activate a neuron is likely to change from one location to the next, and may explain the shift in the cell’s temporal coding properties.

Our data suggests that CA1, the output stage of the hippocampus, does not condense one single representation of “position” for upstream areas, but rather multiplexes multiple transformation of the hippocampal inputs, and autonomously generated representations. This may provide the brain with a palette of computational schemes [60] that may be leveraged to flexibly tackle complex and diverse cognitive tasks.

## Methods

### Subjects

In compliance with Dutch law and institutional regulations, all animal procedures were approved by the Central Commissie Dierproeven (CCD) and conducted in accordance with the Experiments on Animals Act (project number 2016–014 and protocol numbers 0029).

6 male C57BL6/J mice (Charles River) were used in this study, all implanted with a Hybrid Drive. All animals received the implant between 12 and 16 weeks of age. After surgical implantation, mice were individually housed on a 12-h light-dark cycle and tested during the light period. Water and food were available *ad libitum*.

### Surgical Procedures

The fabrication of the Hybrid Drives and the implantation surgeries were done as described earlier [28]. In order to target the dorsal CA1 region of the hippcampus, a craniotomy was made over the right cortex (top-left corner at AP: -1.20 mm; ML: 0.6 mm relative to bregma; bottom-right corner at AP: -2.30 mm; ML: 2.10 mm relative to bregma) using a 0.9 Burr drill. The dura was removed, and the drive’s array was carefully lowered into the brain, with the silicon probe shaft already set to the correct depth. Tetrodes were individually lowered into the brain (5 turns - *≈* 900 *µ*m ). Mice were given at least seven days to recover from surgery before the experiments began.

### Neural and behavioral data collection

Animals were brought to the recording room from post-surgery day 3 and electrophysiological signals were inspected during a rest session in the home cage. Tetrodes were lowered individually, in 45/60*µ*m steps, until common physiological markers for the hippocampus were discernible (SWR complexes during sleep or theta during locomotion). The majority of the tetrodes reached the CA1 pyramidal layer in 7-10 days.

An Open Ephys acquisition board [51] was used to acquire electrophysiological data. Signals were referenced to ground, filtered between 1 and 7500 Hz, multiplexed, and digitized at 30 kHz on the headstages (RHD2132, Intan Technologies, USA). Digital signals were transmitted over two light, custom-made, 12-wire cables (CZ 1187, Cooner Wire, USA) that were counter-balanced with a pulley-system. Waveform extraction and automatic clustering were performed using Dataman (https://github.com/wonkoderverstaendige/dataman) and Klustakwik [29], respectively. Manual waveform curation was performed using the MClust software suit. A CMOS video camera (Flea3 FL3-U3-13S2C-CS, Point Grey Research, Canada; 30 Hz frame rate) was mounted above the linear track and used to record video data.

### Behavioral paradigm

Each behavioral session was preceded and followed by a rest session in the animal’s home cage (“Pre sleep” and “Post sleep”). For the training of the linear track paradigm, mice were positioned at one end of the linear track (1 meter long) with the task to run to the other end to collect a reward (a piece of Weetos chocolate cereal). Another reward was positioned on the opposite end of the trackonly if the animal consumed the current reward. Each lap was defined as an end-to-end run only if the animal’s body started from the first 10 cm of the track and reached the last 10 cm at the other end of the track, without backtracking. Experiments were performed on 10 consecutive days and each behavioral session laster between 20 and 30 minutes. Mice were never food or water deprived.

### Histology

After the final recording day tetrodes were not moved. Animals were administered an overdose of pentobarbital (300mg/ml) before being transcardially perfused with 0.9% saline, followed by 4% paraformaldehyde solution. Brains were extracted and stored in 4% paraformaldehyde for 24 hours. Then, brains were transferred into 30% sucrose solution until sinking. Brains were quickly frozen, cut into coronal sections with a cryostat (30 *µ*m), mounted on glass slides and stained with cresyl violet. The location of the tetrode tips was confirmed from stained sections (Figure S1), in combination with previously mentioned electrophysiological markers.

### Neural data analysis

Before applying other analysis, continuous (LFP) signals were down-sampled to 1 kHz. Most importantly, LFP across the 16 contacts was used to compute the corresponding Current Source Density signal. Since this transformation is based on computing the discrete laplacian along the probe axis (that is along the direction running through CA1 layers), the resulting CSD signal is limited to 14 channels (16 minus the 2 extremes).

### Basins

The CSD signal was used to compute the phase-amplitude coupling strength between a reference theta oscillation and faster oscillations comprised in a [15 250]Hz range. We first applied a wavelet transform to the CSD to obtain the analytical signal over time and across a broad each of these regions, a basin. Small Basins (comprising less than 40 bins in total) were removed from visualizations to improve readability. The same data was then used to track the evolution of the phase-amplitude couplings across different layers and frequencies. For each of the identified main gamma frequency ranges (slow [20–45]Hz; medium [60–90]Hz; fast [120–180]Hz) we extracted layer-specific coupling strength and phad range of frequencies. We then separately obtained the instantaneous phase of the theta oscillation (taken between 6Hz and 10Hz) and the amplitude at the higher frequencies. Theta oscillation reference was generally taken from the stratum lacunosum moleculare, where theta modulation of the LFP is stronger, but the same procedure using a theta from another layer gave equivalent results. Instantaneous theta phase was binned (32 bins) and for each interval the average amplitude of oscillations at higher frequency was computed. We then applied a discrete laplace operator to the frequency-phase matrix so obtained, to identify portions of the matrix characterized by negative curvature. These regions are found by requiring the laplace operator to be negative. For each of these regions, we further selected only the bins that showed a coupling strength above 1STD of the mean of the coupling of the respective frequency. Such operation was repeated for the CSD computed for each contact of the probe. After this double thresholding, each of the layer-specific coupling matrices was then reduced to a sparse version in which only significantly coupled regions had non-zero values. To identify continuities and changes in the couplings across hippocampal layers we then stacked all the sparse-coupling matrices on top of each other, adding a 3rd dimension to the coupling arrangement. Again, we looked for continuous 3D regions of significant bins. We dubbese of the maximum coupling. In the rest of the paper slow gamma power was computed using the signal extracted from the radiatum layer, while medium gamma power was taken from the probe contact in the stratum lacunosum moleculare.

### REM Sleep Detection

Following an established practice [43, 42], periods of REM sleep were classified offline based on the ratio between LFP power in the theta frequency range (5-9Hz) and the power in delta range (2-4Hz). REM periods were identified in the rest period that followed exploration sessions by combining the theta/delta ratio with an additional requirement for absence of animal movement.

### Gamma Coefficient

The instantaneous balance between the power in the slow gamma (*P_Slow_*(*t*)) and medium gamma (*P_Med_*(*t*)) frequency range was computed as follow. First both the slow gamma power and medium gamma power during running periods were separately z-scored. Then a power-ratio score, spanning the [-1 1] interval was computed as:

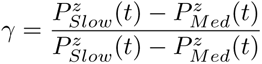

so that a value of -1 would correspond to total medium gamma domination and +1 to slow gamma completely dominating.

### Place Cell Identification

Putative excitatory pyramidal cells were discriminated using their auto-correlograms, firing rates and waveform information. Specifically, pyramidal cells were classified as such if they had a mean firing rate < 8 Hz and the average first moment of the autocorrelogram (i.e., the mean value) occurring before 8 milliseconds [12]. Only cells classified as pyramidal cells were used for further place cell analysis. Place cells were defined applying a combination of different criteria. All analysis were performed on speed-filtered activity, after removing periods in which the animal speed was smaller than 5cm/s. Then, only cells with an average activity above 0.3 Hz were taken into account. Then for each of these cells, the Skaggs information per second 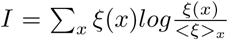 was compared to distribution of information values resulting from randomly shuffling cell spike times. A cell passed the selection if its Skaggs information was significantly higher than the ones of the surrogate distribution. Last, only place cells with peak firing rate higher than 1 Hz were kept for further analysis. Place fields were isolated as continuous regions in the rate map with rate higher than 20% of the cell maximum activity. Multiple fields (field one and field two) were then sorted according to their respective peak firing rate.

The modulation of place field shape by gamma power was evaluated by first estimating the place field boundaries using all spikes from a cell. Then the place field extension was divided in 7 spatial bins. For each of these bins we computed the spike rate of the cell using only spikes emitted in association of a specific range of the slow-to-medium gamma power ratio. For each cell the overall spike probability over the bins was normalized to 1 and the final value for each bin was obtained as an average over all cells in the dataset. Opposite running directions were treated separately.

### Place Cell Classification

Spikes were filtered both in speed (>5cm/s) and in position, by taking only those emitted within the boundaries of the cell place field (field one in the case of multiple fields). For each place cell, we measured to what degree the phase (with respect to theta oscillations) of spikes was concentrated and to what extent it was modulated by the respective position within the place field. In the first case we computed the length of the Reyleigh vector associated to the distribution of spike phases with respect to the instantaneous theta phase (slm theta). In the second case we used a circular-linear correlation to measure the degree of dependency between the spike phase and the position within the field. Both measure yielded a score between 0 and 1, with 1 meaning perfect phase locking and perfect phase-position relation, respectively. We combined the two scores into a ’Theta Score’ subtracting one from the other: Theta Score = Precession Score - Locking Score. Because of the similar interval of the two measures, negative Theta Scores would indicate spikes probability being mostly modulated by specific phases of theta, while positive Theta Scores would point to a substantial presence of phase precession.

### Single Spikes and Burst Spikes classification

We subdivided spikes from a cell into single and bursts by applying a temporal interval filter. Any spike occurring within 7ms of another was labelled as belonging to a burst, all the others were considered as single spikes. If not otherwise stated, spike probability evaluation and other analysis were performed using only single spikes. Burst spikes were analyzed separately.

Significance of theta phase selectivity for either single or burst spikes was evaluated with a shuffling procedure. The probability of spikes in any bin along the [0 2*π*] interval was computed by dividing the number of spikes of each cell by the probability of observing a specific gamma coefficient in the same theta phase interval. For each cell, this ratio was compared to an equivalent measure applied to a surrogate version of the cell activity, where spike times had been randomly reassigned to any of the temporal bins characterized by a slow-to-medium gamma power ratio in a specific range. Such comparison was repeated 1000 times for each cell, each time repeating the shuffling procedure. Statistics were computed on the distribution of differences between the real spike distribution and the surrogate ones.

### Generalized-Linear Model (GLM)

As described in previous work [19] we used a maximum entropy model inference paradigm to reconstruct the distribution of each cell’s firing probability. As a statistical model, we considered the maximum entropy model known as kinetic Ising model. We first separated running periods from periods of quiescence by applying a 5 cm/s speed filter. The activity of the cells was binned in 10 ms bins, and a binary variable *S_i_*(*t*) was assigned to each neuron for each temporal bin. *S_i_*(*t*) was taken to be a binary variable, with +1/*−*1 values, depending on the presence/absence of spikes emitted by neuron i within time bin t. Since the length of time bins was relatively short, this was a reasonable approximation as the case of multiple spikes per bin was rare. Letting the state of each neuron at time t depend on the state of the population in the previous time step t *−* 1 , the maximum entropy distribution over the state *S_i_*(*t*) of neuron i at time t is

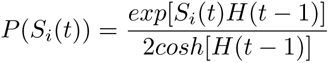

where *H*(*t*) is a time dependant covariate having the role of the external field in statistical physics. Eq. 1 defines a GLM (Generalized Linear Model), where, in each time bin, mostly only one or zero spikes per bin are observed and the interaction kernel extends one time step in the past. To find what values of *H*(*t*) are the most likely to generate the observed data given Eq. 1, we maximized the log-likelihood function

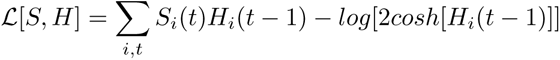

with respect to *H*(*t*). The log-likelihood measures how well the model explains the statistics in the observed data. In our analysis, we have used the natural logarithm. Since the external field, *H*(*t*), can explain the variations in the firing rate as the rat navigates in space, it becomes important to model it appropriately. Here we assumed that the spatial input arises as the sum of one-dimensional Gaussian basis functions centered on evenly spaced L intervals covering the linear track. Opposite running directions were treated as distinct environments and analyzed separately. The spatial field of cell i at time t is then

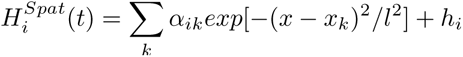

where *h_i_*is a unit-specific spatially and temporally constant baseline, and *x_k_* and *l* are the centers of the regular intervals and the widths of the basis functions, respectively. We expanded this purely spatial description of cell activity by combining the spatial contribution to activity with that of specific phases of theta oscillation and a specific slow-medium gamma balance. To do so, we tiled the 3-dimensional space obtained from combining i) the position on the track, ii) the theta phase (in the [0 2*π*] interval) and the relative strength of the instantaneous power in the slow and medium gamma range, computed as a normalized difference: 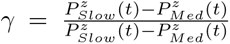 (spanning the [-1 1] interval), where the *z* apex indicates previous z-scoring of both power time-series. Such space was tiled with a regular square lattice of L x T x G three-dimensional Gaussians, with 0 off-diagonal terms and variance *l*, *t* and *g* in the three directions. The external field of cell *i* can thus be expressed as:

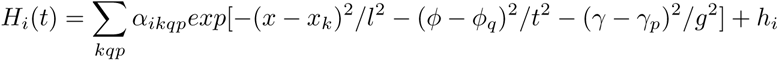

where we introduced the dependency on the current phase *φ* of the theta oscillation and current gamma balance *γ*. An accurate representation of the cell activity in the space-phase configuration, can be found by inferring the parameters *α_ikqp_* of the linear combination of the Gaussian basis function. We first optimized the values of L and T (the number of Gaussian basis functions in the lattice) and l, t (their widths), while keeping G and g fixed (G = 6 and g=0.4). We maximized the likelihood over a range of values of L (from 15 to 25), T (from 4 to 10), l (from 5 to 30 cm) and t (from 0 to *π/*2) and chose the values of the parameters that gave the highest Akaike-adjusted likelihood value. The Akaike information criterion (AIC) is a measure to compensate for overfitting by models with more parameters, where the preferred model is that with the minimum AIC value, defined as

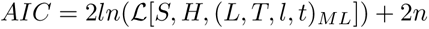

where *L* is the likelihood at the maximum likelihood (ML) estimates of the parameters *α*’s and *h_i_*) for a given value of M, N, r and p. *n* is the number of parameters (here r and p do not affect this number as it is a scale factor for Gaussian basis functions, while larger values of M and N result in more parameters *α* included in the model). The procedure was performed over all available sessions at once so that the resulting optimal parameters (L=20, T=6 and l=8 cm, p = *π/*3) were applied to all of them. Firing rate maps can then be expressed for an arbitrary combination of position, phase and gamma balance as

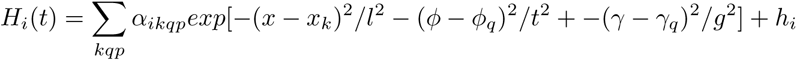

where *x*, *φ* and *γ* are the desired values of position, theta phase and gamma balance. In all our analysis, we considered a partition of the environment in bins of 2.5cm. The effects of speed on the firing probabilities was addressed in a similar manner, by including the instantaneous velocity of the animal as a further covariate in the Gaussian decomposition method. The GLM was then ran to infer a set of *α_ikqpz_* weights covering a 4D space of position x phase x gamma x speed. The parameters of the Gaussian-basis along the speed dimension were not optimized but were taken as *V* = 4 Gaussians with a standard deviation of 6cm/s.

### Pairwise Spike Timing

For each cell, spikes emitted during running were further filtered so that: i) only first spikes in each theta cycle (taking the peak of the theta oscillation as the starting point of the cycle) and ii) only spikes emitted within the cell place field one, were considered. For each of these spikes we also computed the simultaneous slow/medium gamma power ratio which we then used as a label to further subdivide the spikes. We then considered all the cell pairs in the population of simultaneously recorded place cells. Each pair (A,B) of place cells was then defined by i) the spatial distance between the place fields of cell A and B (computed as the distance between the centers of mass, but using the distance between the fields peaks did not change the results) and ii) the average phase interval between the spikes of the two cells. The latter was computed by taking spikes of cell A as reference and within each theta cycle (for which a spike from both cells was available) taking the *dθ_A−>B_*(*k*) (where k stands for the kth theta cycle), that is the (signed) difference between the phase of the spike of cell A and that of cell B spike. We then computed the center of mass of the *dθ_A−>B_* distribution and used it as a measure of the average phase offset between that cell pair. Spatial distance and phase distance were then arranged in a 2-dimensional space and the average phase offset was computed over different ranges of spatial distances as long as at least 10 pairs of cells were available. The analysis was repeated using cell pairs from specific populations subgroups and by restricting the use of spikes to those emitted in the presence of a certain gamma balance.

### Bayesian decoding

To reconstruct the position on the track encoded in theta associated spiking activity, we implemented a standard Bayesian decoding procedure [63]. Periods of active locomotion were first segmented using the peaks of theta oscillation (extracted from the stratum lacunosum moleculare). Each theta period was further subdivided using a sliding window of 30ms length and with an offset of 10ms. A population vector was then built for each of these sub-windows ***η***(*t*), where t indicates the time within the theta cycle subdivision. The likelihood *P* (*x|****η***(*t*)) of position on the track *x* given the population activity was then computed according to Bayes’ formula using the set of probabilities *P* (***η***(*t*))*|x*) obtained from marginalizing the result of the GLM (see above) over the phase of spiking. The resulting probability density *P* (*x, t*) was normalized so that ∑*_x_ P* (*x, t*) = 1 for each time window *t* that contained 3 spikes or more, and was otherwise set to 0 and ignored in the following analysis. Theta cycles with less than *n* ’active’ time windows were removed from the following analysis (where *n* was set to 5 when considering the entire cell population and lowered to 3 when decoding from subgroups of cells). Using the normalized probability density, for each theta cycle we computed two scores. We first evaluated the amount of ’local’ information in the activity by summing the probability concentrated around the current position of the animal ϒ*_Local_* = ∑*_t_ P* (*x^∗^, t*) where *x^∗^* denotes the set of spatial bins in a range of 5cm of the animal position at time *t*. The presence of ’non-local’ sequence-like activity spanning a significant spatial interval around the animal position was instead evaluated by running a set of linear regression over a modified version of the *P^∗^*(*x, t*) matrix where the ’local’ decoding probability density had been subtracted out of the original *P* (*x, t*) (so that we could factor out the effect of positional decoding from the reconstructed trajectory). Among these regressions, we searched for the best-fit, overlapping with the largest amount of probability density. That is, Υ*_NonLocal_* = *max_β_* ∑*_k_ P^∗^*(*x_β_*_(_*_k_*_)_*, t_β_*_(_*_k_*_)_) where *β* indicates a set of linear parametrizations of *x* and *t* : *x* = *β*_1_ + *t × β*_2_. Importantly *β*_2_ was taken to be always *>* 0 so to exclude fits very close to the =ϒ*_Local_* defined above. The same procedure was applied using different set of cells to compute the probability density, so to obtain an estimation of the spatial information carried by specific cell groups (in our case, the entire population and either phase precessing or phase locking cells). Theta cycles were further classified according to the simultaneously expressed gamma power ratio. The average ratio between the power of slow and medium gamma over the interval of the theta oscillation was used as a label for that theta cycle and used to subdivide the set of cycles depending on the relative strength of the slow and medium gamma component. These scores were used to evaluate the content of activity in different theta cycles. An additional step was then taken to compare the nature of activity across different cell populations. In this case the decoding scores obtained from each cell group were first normalized to *max*_Θ_ϒ = 1 (where the max is taken over all available theta cycles). For each theta cycle in which enough spikes were emitted by cells from both independent cell groups we computed the difference between the two obtained decoding scores ϒ_1_ and ϒ_2_ (either Local or NonLocal) as the distance of the point (ϒ_1_,ϒ_2_) from the equal-score diagonal ϒ_1_ = ϒ_2_.

## Statistics

Data analyses were performed using custom MATLAB scripts (The Math Works). Paired and unpaired t-tests, Kolmogorov-Smirnov tests were performed using standard built-in MATLAB functions. 2-D Kolmogorov-Smirnov test was performed using custom code based on the Peacock algorithm. All tests were two-tailed, except where stated otherwise. Linear regressions and confidence intervals over the fit parameters were obtained using standard MATLAB functions (i.e. ’fit’ and ’confint’). The significance of the difference between two fit slopes was estimated using the degree of overlap between the two confidence intervals. For permutation tests, independent variables were shuffled 1000 times and this null distribution was compared to the one obtained from data.

## Data and Software Availability

All computer code and all data will be made available upon a reasonable request to the Lead Contact, Francesco P. Battaglia.

## Acknowledgments

We thank Freyja Olafsdottir, Cliff Kentros and Alessandro Treves for their comments on the manuscript and the Battaglia lab for helpful discussions and comments at different stages of the project. This work was supported by the European Union’s Horizon 2020 research and innovation program (MGate, grant agreement no. 765549; M.G. and F.P.B.), the European Research Council (ERC) Advanced Grant “REPLAY-DMN” (grant agreement no. 833964; F.P.B.), European Union’s Horizon 2020 Research and Innovation Programme Grant “BrownianReactivation” (grant agreement no. 840704; F.S.).

## Author contributions

Conceptualization: M.G., F.S. and F.P.B.; Investigation and Data Curation: M.G.; Methodology and Software: M.G, F.S. and F.P.B.; Resources: F.S. and F.P.B.; Formal Analysis and Visualization, F.S. and M.G.; Writing – Original Draft: M.G., F.S. and F.P.B.; Writing – Review & Editing: M.G., F.S. and F.P.B.; Supervision and Funding Acquisition: F.S. and F.P.B.

## Extended Data

**Figure S1:**
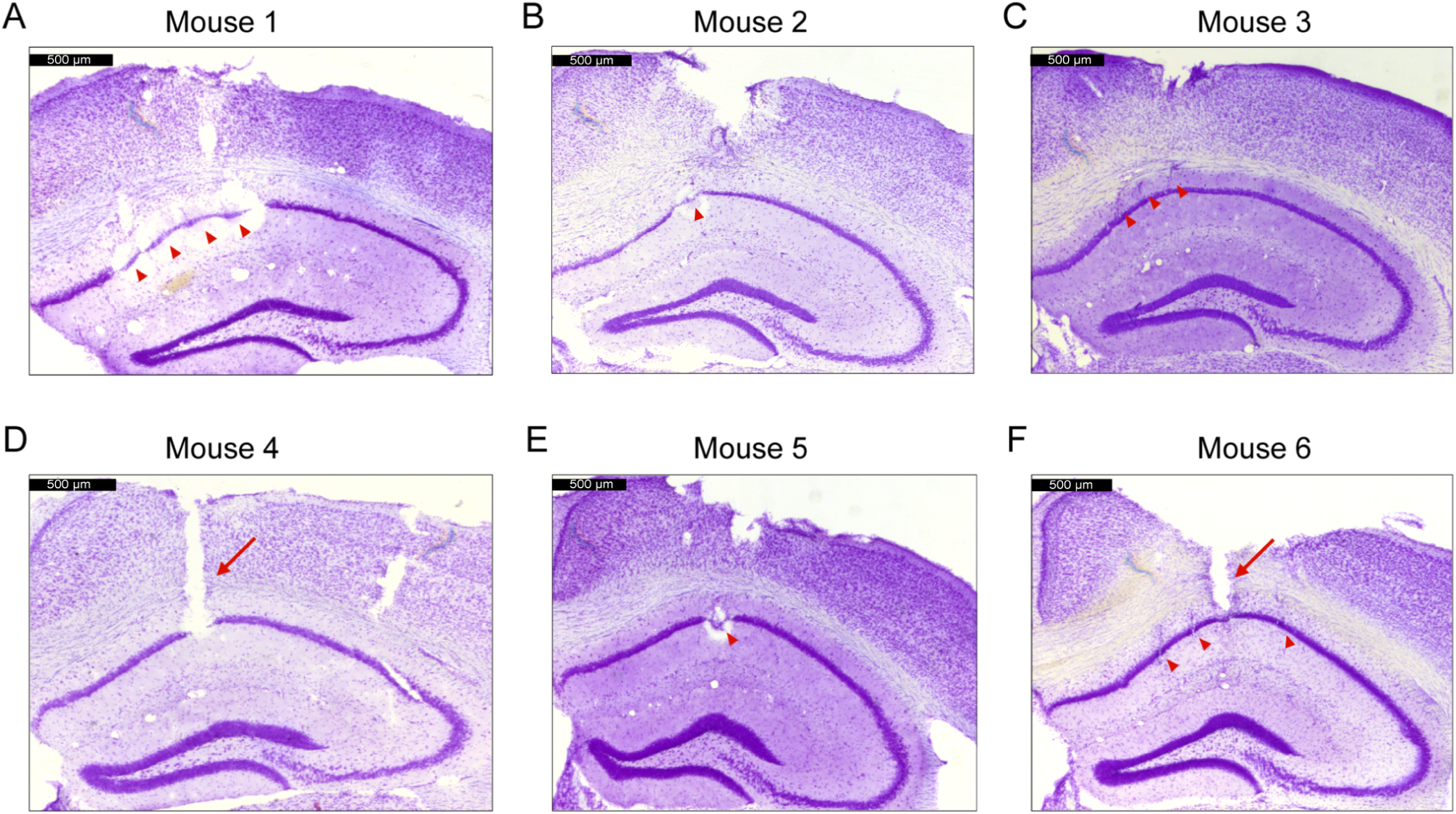
Histological verification of tetrodes and silicon probe position. Representative Nissl-stained coronal section showing example recording locations from each animal. Red triangles indicate the estimated location of tetrode tips (CA1 pyramidal layer), while red arrows indicate the trace of the silicon probe shaft. Mouse 1, 2 and 5 received electrolytic lesions at the end of the experiment. Mouse 3, 4 and 6 never electrolytic lesions, to better visualize the silicon probe track. Scale bar denote 0.5 mm.

**Figure S2:**
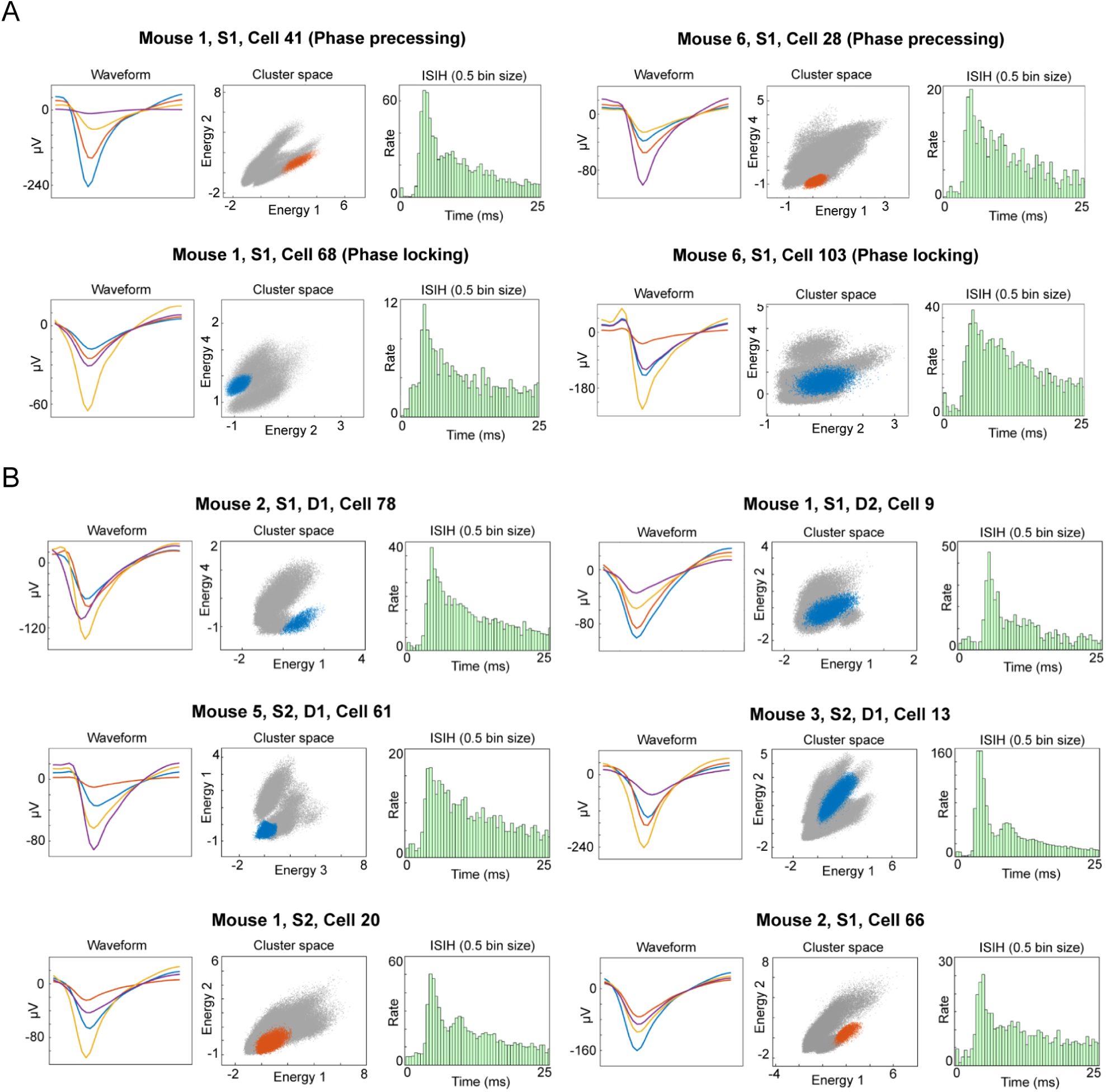
Cluster information of individual single unit examples. Left panel, mean waveform across the 4 channels of the tetrode. Center panel, 2D cluster projection of the recorded spikes. The cell’s cluster is highlighted by blue or orange color. Right panel, interspike-interval histogram (0.5 ms time bins). (A) Information for the individual units reported in Figure 2C. (B) Information for the individual units reported in Figure 3C,F.

**Figure S3:**
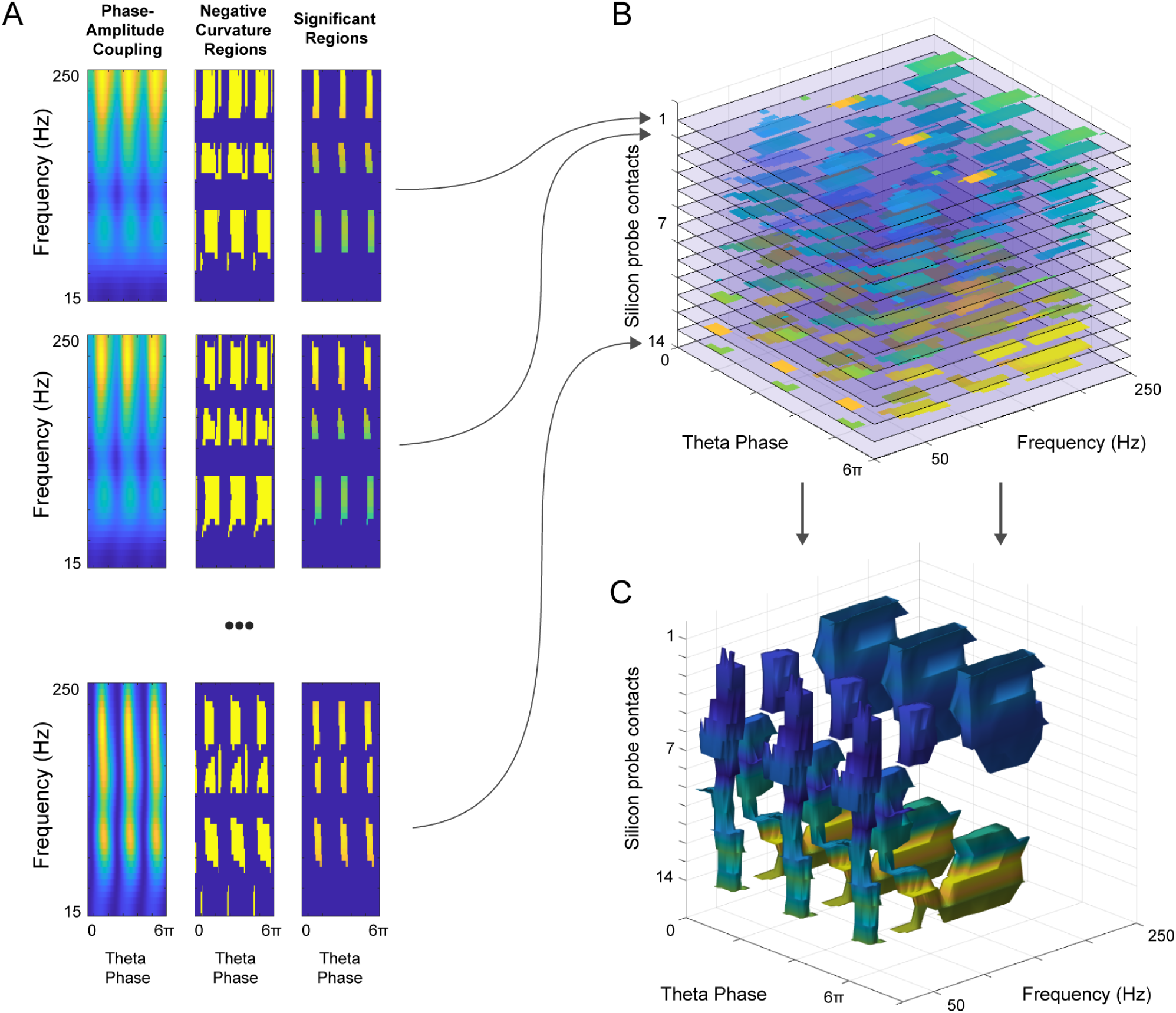
Construction of a 3D phase-amplitude-depth structure to visualize oscillatory interactions at different frequencies. (A) Left panel, phase-amplitude plot between the theta oscillation (6-10 Hz) and the power of oscillations at higher frequencies (15-250 Hz). Center panel, negative curvature regions found applying a discrete laplace operator to the frequency-phase matrix obtained in left panel. Right panel, matrix with further selected bins that showed a coupling strength above 1STD of the mean of the coupling of the respective frequency. Each analysis was repeated for each CSD-derived signal from 14 contacts of the silicon probe (16 minus the 2 extremes). (B) Stacked matrices with bins that showed a coupling strength above 1STD of the mean of the coupling of the respective frequency (same as in right panel (A)). (C) 3D Basins structure obtained concatenating the significant coupling values in Panel (B) at different depths.

**Figure S4:**
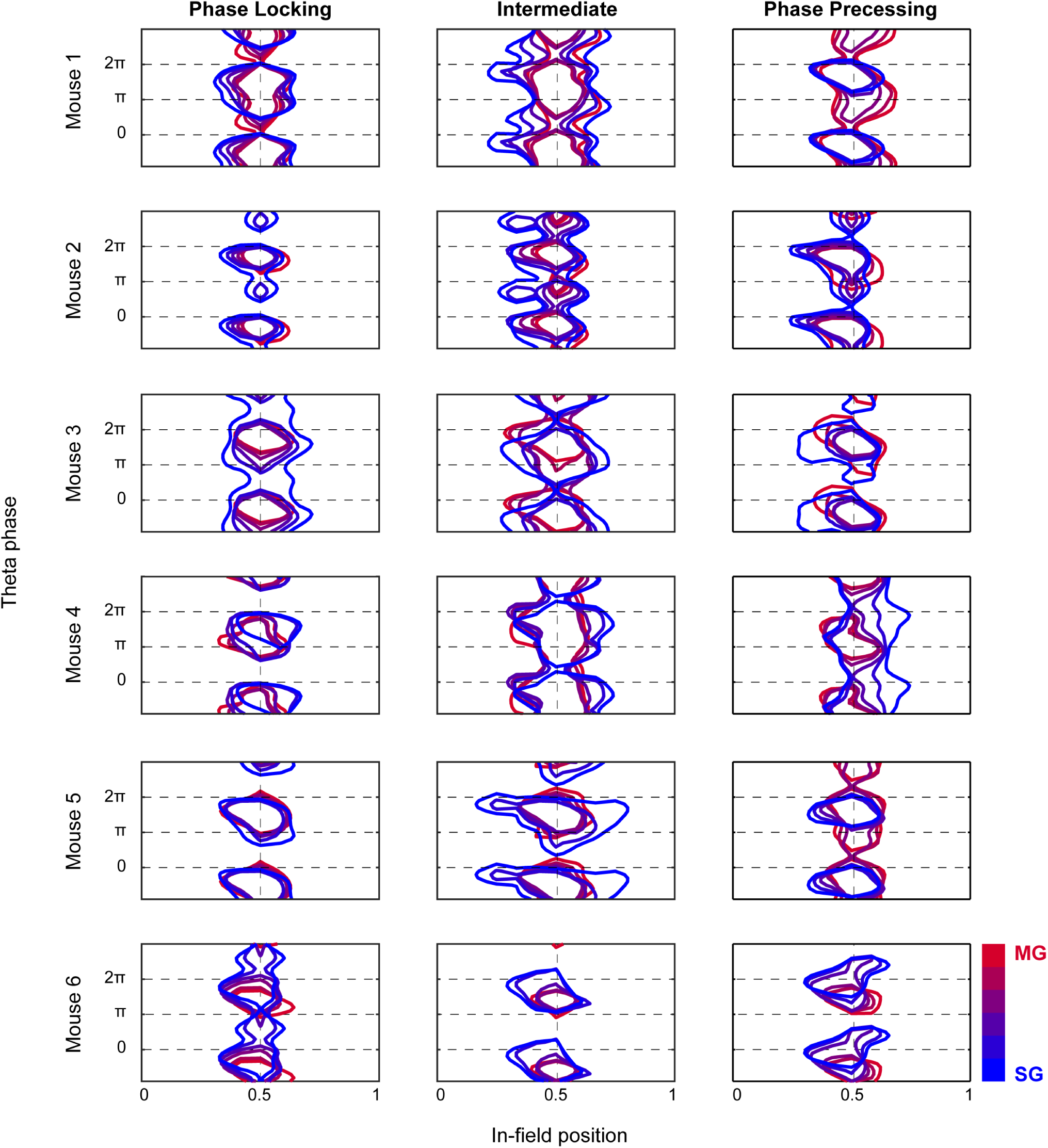
Spike density contour plot for each animal. Contour plot of the GLM-derived spike density plot (same as in 5B) for each animal, as a function of the slow-medium gamma coefficient.

**Figure S5:**
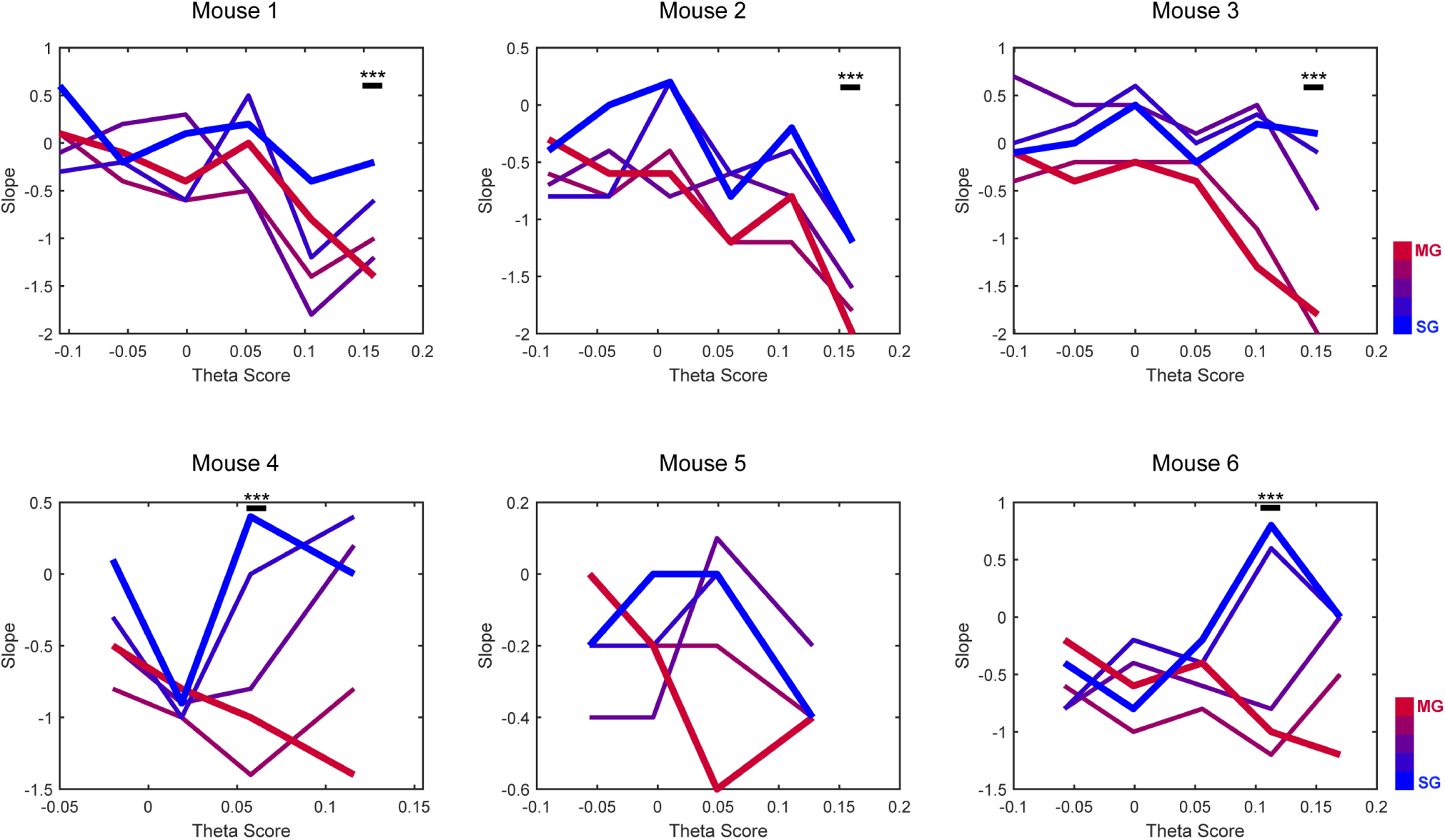
Single cell regression plots for each animal. Phase-position slope values plotted against the Theta Score values for each individual field, as a function of slow-medium gamma balance (all animals). p<0.05; Spearman Correlation.

**Figure S6:**
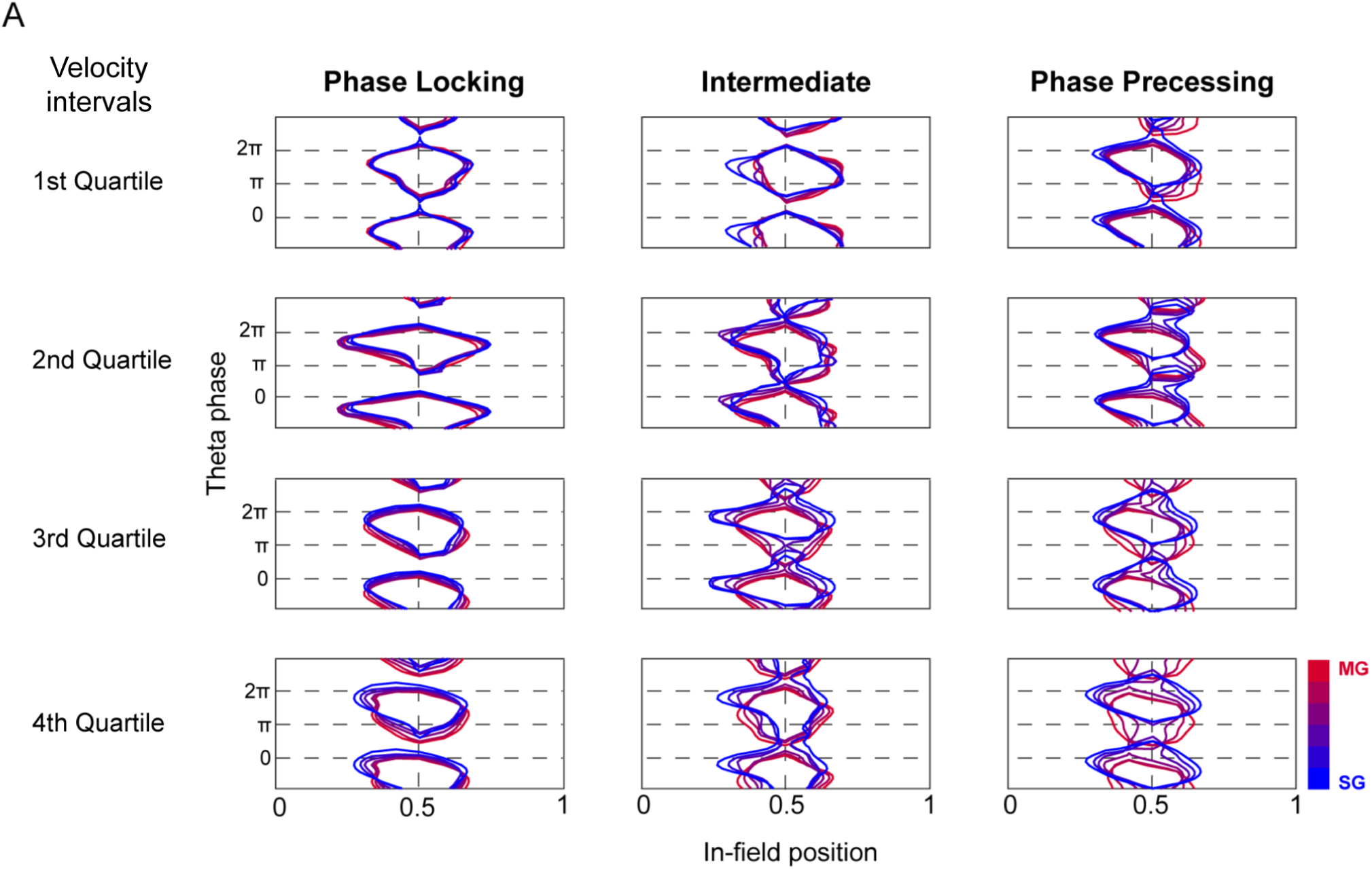
Spike density contour plot using velocity as an additional factor in the GLM. Contour plot of the GLM-derived spike density plot as a function of velocity. Same amount of data in each velocity-interval group. All animals (N=6), all sessions (n=12).

**Figure S7:**
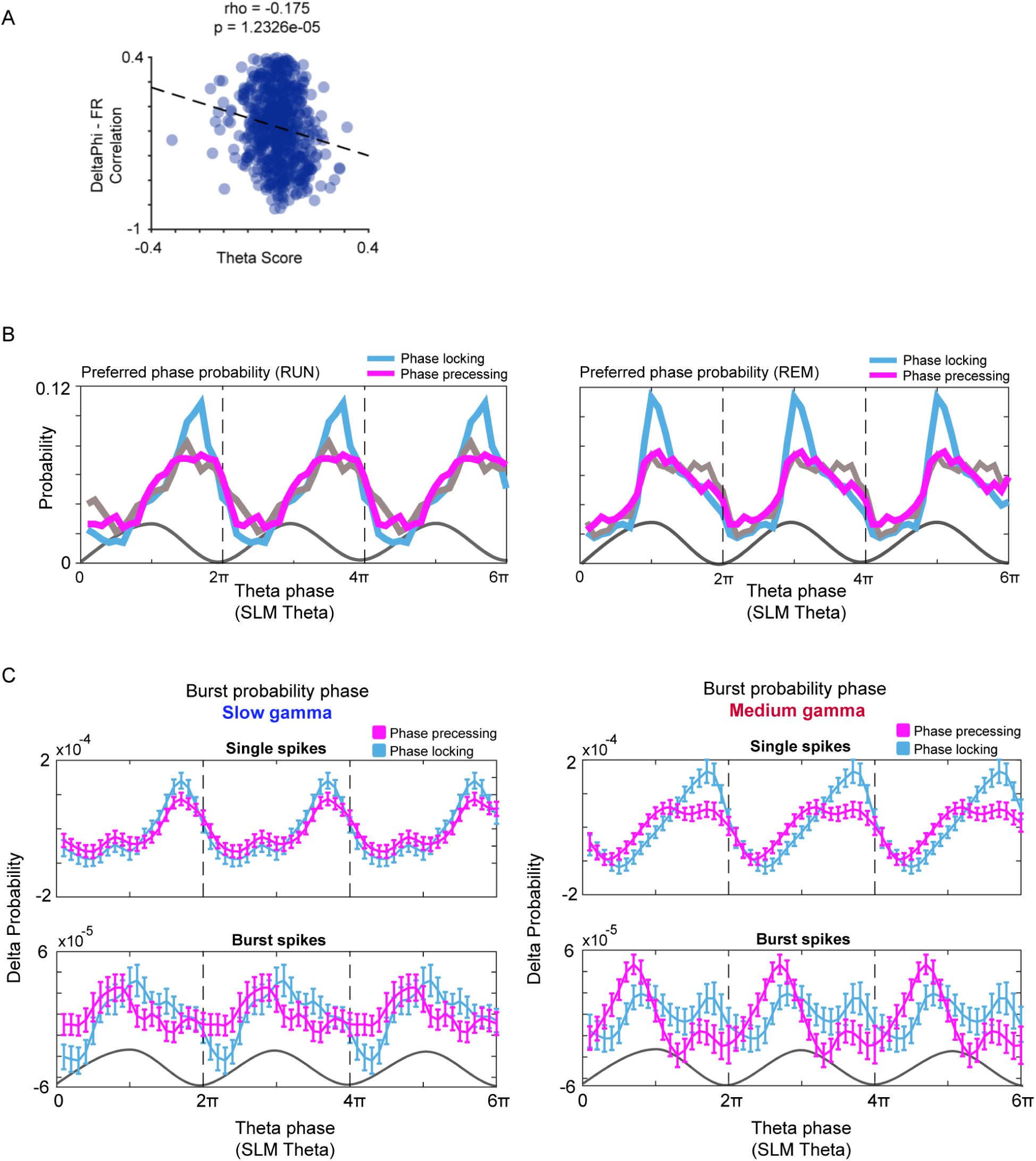
Additional spike-phase relation properties. (A) Correlation between the Theta Score and the DeltaPhi - Firing Rate (FR) correlation for field one of each cell (p<0.01; Spearman’s Rho = -0.175). (B) Preferred firing phase preferences for phase precessing, intermediate and phase locking cells during REM sleep. (C) Phase distributions of single spikes and burst spikes during slow (left) and medium (right) gamma dominated periods, for phase locking (light blue) and phase precessing (violet) cells. Rayleigh Vector Length Difference of the peaks against shuffling is significant (T-test; p<0.01) (D) Phase distributions of single spikes and burst spikes during medium gamma dominated periods, for phase locking (light blue) and phase precessing (violet) cells.

**Figure S8:**
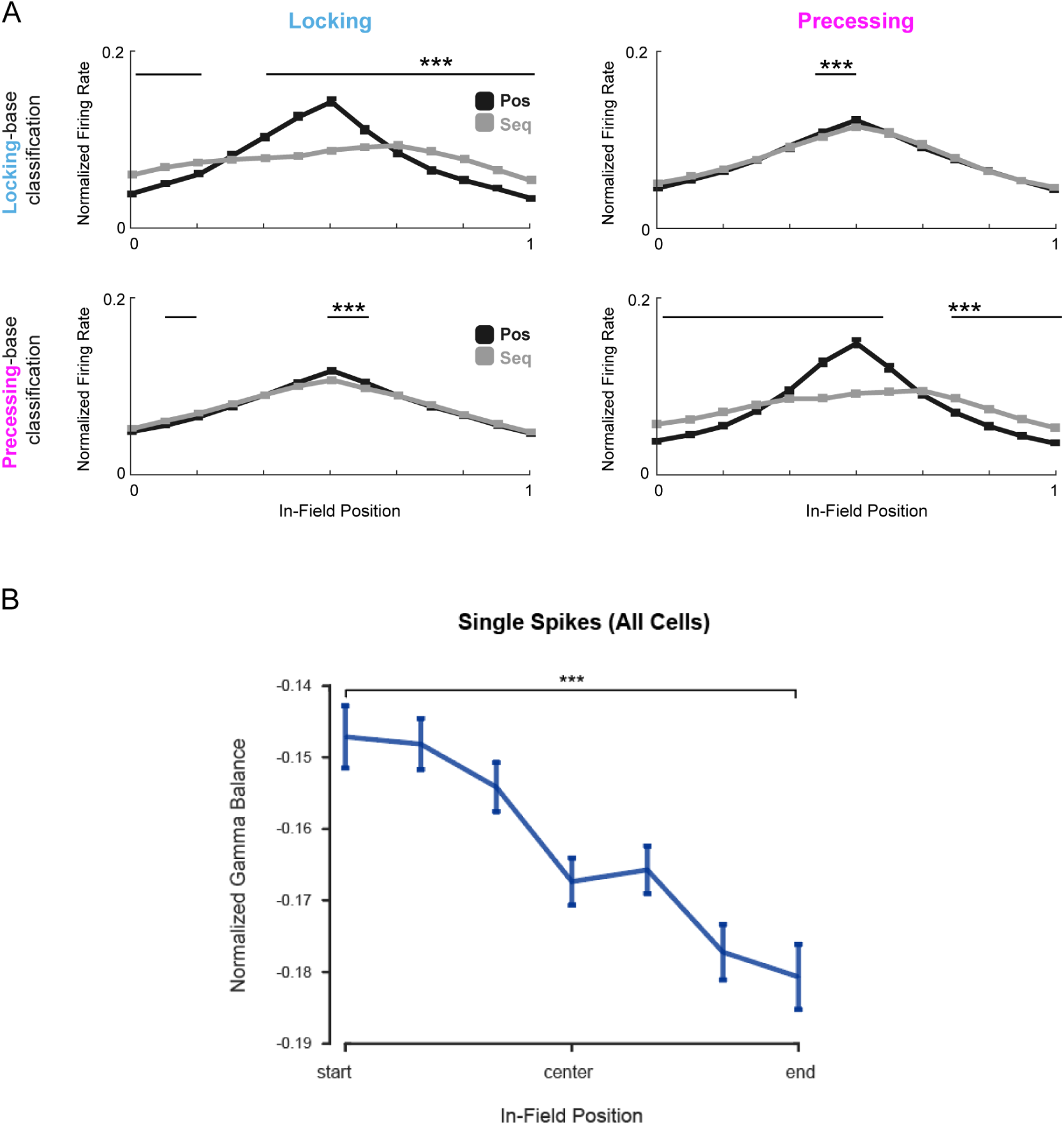
In-field dynamics as a function of place field populations and gamma coefficient. (A) Place field shape obtained when considering different temporal intervals defined by the type of spatial coding observed, by selecting theta cycles based on the position vs. sequential decoding score. Locking vs precessing fields. Top: classification based on phase locking fields. Bottom: classification based on phase precessing fields. (p<0.01; t-test). (C) Instantaneous medium/slow gamma balance as a function of in field position. Normalized gamma balance (medium gamma vs slow gamma) within the cell’s place field. Values at the start and the end of the place field are statistically different (p<0.01; two-sample t-test).

